# Data-Driven Mechanistic Computational Modeling of Human Cell Cycle Phases for Defining Inhibitor Combinations in Silico

**DOI:** 10.64898/2026.02.04.703693

**Authors:** Justin A. Womack, Andrew Sukowaty, Ansel J. Fellman, Ranjan K. Dash, Scott S. Terhune

## Abstract

The human cell cycle is a highly regulated process that integrates multiple signaling pathways and checkpoints to ensure faithful genome duplication and cell division. Disruptions in these regulatory networks contribute to a wide range of diseases. Here, we present a novel, updateable computational model of the full human cell cycle that shows sustained oscillations over time and reproduces experimental perturbations. We used a hybrid framework combining mass action and Michaelis–Menten kinetics, incorporating the synthesis, degradation, and regulation of key cell cycle proteins and protein complexes. It consists of 63 distinct biochemical species, interacting through 41 major reactions, and functioning through 63 ODEs. The model is built upon a modular framework, structured around the core regulatory networks of the G1, S, G2, and M phases. Due to its complexity, we determined parameter sets that met strict criteria, namely event timing, comparable concentrations, and continuous cycling. We validated the model’s behavior by reproducing canonical checkpoint responses, including mitogen dependence and the DNA damage response, both of which produced reversible and robust cell cycle arrests. Importantly, the model was trained and calibrated using *in vitro* data from human U251-MG glioma cells expressing the FastFUCCI cell cycle reporter. We quantitatively aligned the simulated and experimentally determined phase durations and cell doubling times. Next, we experimentally tested and refined model parameters by using abemaciclib-mediated inhibition of CDK4 and volasertib-mediated inhibition of PLK1. *In vitro* and *in silico* data show dose-dependent G1 arrest by abemaciclib and dose-dependent mitotic arrest by volasertib. Finally, we demonstrated that the model predicts changes in cell proliferation over a wide range of drug concentrations and combinations. Overall, our work establishes a robust, data-driven computational model for systems-level analysis of the human cell cycle and its disruption by therapeutic perturbations.

**AUTHOR SUMMARY:** Knowledge of the protein-protein interaction networks governing the cell cycle is ever-expanding, yet this information is often fragmented across studies focusing on disconnected subsets of the cycle. For decades, researchers have investigated the underlying mechanisms of cell division, but an integrated, quantitative understanding of the entire process remains elusive. This gap is a major hurdle for predicting how targeted therapies affect cell proliferation, especially when used in combination. Our goal is to develop an *in silico* simulation of the complete human cell cycle by integrating the key mechanistic relationships across all four phases into a single computational model with enough resolution to approximate outcomes upon perturbation. In achieving this, we have developed a novel, comprehensive computational model that provides an integrated quantitative understanding of how cancer drugs alter the human cell cycle. We have rigorously trained, calibrated, and validated the model by suitably estimating its parameter values to produce accurate cell cycle phase timing, using high-resolution, live-cell imaging data and other cardinal features of the cell cycle in a U251-MG glioblastoma line. This work provides an accessible tool for exploring how normal cell cycle control is disrupted in disease, generating new hypotheses, and identifying potential points of therapeutic intervention.

## INTRODUCTION

Progression through the mammalian cell cycle involves a coordinated sequence of growth and division events that result in the formation of two daughter cells. This process is classically divided into four phases: Gap 1 (G1), DNA synthesis (S), Gap 2 (G2), and Mitosis (M) phases [1, 2]. Strict regulatory mechanisms ensure that the genome is duplicated once per cycle and accurately segregated, preserving genomic integrity. Disruption of these controls can lead to aneuploidy, aberrant cell death, or uncontrolled proliferation, hallmarks of numerous human diseases, including cancer [3–7]. Cells may exit the active cycle into a quiescent G0 state, where most terminally differentiated cells reside [8, 9]. In contrast, proliferative signals such as growth factors, cytokines, or receptor engagement can stimulate cell cycle re-entry. Upon entering G1, cells express factors required for DNA replication, followed by genome duplication during S phase, quality control and preparation in G2, and mechanical segregation of sister chromatids during M phase [10]. Progression through these phases depends on tightly regulated temporal changes in protein expression, activity, post-translational modification, subcellular localization, and protein–protein interactions [11–13]. Together, these coordinated processes ensure robust and ordered cell cycle progression.

The cell cycle is governed by feedback-controlled molecular switches built around cyclin-dependent kinases (CDKs) and their cyclin partners [14]. These switches generate bistable states that enable decisive, irreversible phase transitions while maintaining responsiveness to perturbations such as DNA damage or growth factor withdrawal [15, 16]. CDK activity is regulated through three principal mechanisms: (1) oscillatory cyclin synthesis and ubiquitin-mediated degradation [17], (2) reversible phosphorylation by CDK-activating and inhibitory kinases and phosphatases [18], and (3) inhibition by CDK inhibitor (CKI) proteins [19, 20]. Cyclin levels are regulated transcriptionally by E2F family transcription factors [21–24] and post-translationally by ubiquitin-mediated degradation through the Anaphase-Promoting Complex/Cyclosome (APC/C) [25–31] and Skp1–Cullin–F-box (SCF) E3 ligase complexes [17, 32–34], forming layered control that enforces ordered, irreversible cell cycle progression.

Given the complexity and tight coupling of these regulatory processes, experimental approaches alone often struggle to identify key system-level mechanisms. This challenge is especially pronounced in the development of cancer therapies, where the combinatorial space of potential treatments far exceeds what can be experimentally tested. As a result, integrated computational modeling has emerged as a powerful approach for simulating dynamic cellular behavior, generating testable hypotheses, and predicting responses to therapeutic interventions *in silico*. Recognizing its potential, the U.S. National Institutes of Health (NIH) has identified the integration of computational modeling with experimental biology as a major research priority for understanding disease mechanisms and identifying therapeutics [35].

Ordinary differential equation (ODE)–based models have been widely used to translate biochemical reaction networks into mathematical frameworks capable of simulating dynamic cell cycle behavior (Reviewed in [36]). Despite decades of progress, a comprehensive, experimentally grounded computational model of the complete mammalian cell cycle remains elusive. Early models established the qualitative mathematical foundations of cell cycle oscillations but relied largely on parameter tuning rather than quantitative experimental data [37–40]. Subsequent mammalian cell cycle models expanded biological coverage to include additional checkpoints, growth factor signaling, and DNA damage responses at G1/S and G2/M transitions. However, many of these models continue to exhibit persistent limitations, including fragmentation across cell cycle phases, limited quantitative validation, and an inability to sustain stable oscillations without external resetting [38, 41–45]. For example, Weis et al. [46] combined flow cytometry data with an ODE-based model of key cell cycle regulators, achieving agreement with experimental measurements across multiple time points in a human cell line. Despite this quantitative fitting, sustained oscillations required manual resetting of select G1 variables. More recent efforts have begun integrating experimental measurements and large-scale parameter optimization [47], yet challenges remain in achieving predictive, experimentally validated, and fully oscillatory representations of the complete mammalian cell cycle.

To address these challenges, we constructed a data-driven, mechanistic computational model of the complete human cell cycle. The initial framework combined the skeletal architecture developed by Gerard et *al.* [48, 49] with our previously validated mitosis models [50, 51]. We extended the regulatory pathways governing G1, S, and G2 phases while preserving the core mitotic dynamics. This produced a cohesive system capable of simulating both normal and dysregulated cell cycle behavior. We calibrated and validated the model against high-resolution data from a U251-MG glioma cell line using the FastFUCCI system (Fluorescent Ubiquitination-based Cell Cycle Indicator) [52]. Together, this work establishes a holistic, experimentally grounded computational framework for investigating human cell cycle regulation and therapeutic perturbations.

## RESULTS

### Formulation of a modular and expandable framework for the human cell cycle, including cyclins and their regulators

We have developed a unified computational model of the human cell cycle as a system of ordinary differential equations (ODEs). This model consists of 63 distinct biochemical species (**Appendix S1**), interacting through 41 major reactions (**Appendix S2**) and functioning through 63 ODEs (**Appendix S3**). The model is built upon a modular framework, structured around the core regulatory networks of the G1, S, G2, and M phases, with each phase transition driven by the precisely timed activity of a dominant Cyclin-CDK complex. The skeleton of this model includes concepts introduced by Gérard et al. [10]. Their model integrated the regulatory modules driving G1, S, G2, and M phases into an oscillatory system. In our updated model, the underlying network logic uses a hybrid kinetic framework that combines mass-action kinetics for association and dissociation with Michaelis-Menten kinetics for enzymatic processes such as synthesis and degradation. The model also incorporates our previously developed framework regulating mitosis [51, 53]. This overall structure allows the model to capture the complex interplay of feedback loops and bistable switches that govern irreversible cell cycle progression.

The mammalian cell cycle is organized around several core Cyclin:CDK complexes that act as sequential, phase-specific drivers of progression (**Fig. 1A**). The complexes that we account for include Cyclin D (CCND) binding CDK4 or 6 (4/6) for G1, Cyclin E (CCNE) binding CDK2 for G1/S, Cyclin A (CCNA) binding CDK2 for S/G2, and Cyclin B (CCNB) binding CDK1 for G2/M. Several of these factors have multiple cell-type-specific variants, which we do not account for in this initial base framework. Each complex rises and falls in both abundance and activity, regulated by diverse signals. For simplicity, the regulatory signals are numbered S1- S11 (**Fig. 1**). In G1, the CCND:CDK4/6 complex (**Fig. 1B**) is the first to rise in response to extracellular mitogenic stimulation, with signaling inducing CCND synthesis [54–56]. CCND binds CDK4/6 to form an active kinase that phosphorylates RB1 and initiates G1 progression [57, 58]. Its activity is further fine-tuned through reversible phosphorylation: WEE1 kinase-mediated phosphorylation inactivates the complex while CDC25A phosphatase removes this inhibitory signal [59]. Eventually, CCND is marked for degradation by SCF-mediated polyubiquitination, providing negative feedback that limits its duration and ensures proper handoff to the next phase [57, 60]. Our model degrades cyclins that are both free and complexed with their respective CDK. The G1/S transition is driven by the CCNE:CDK2 complex (**Fig. 1C**). CCNE synthesis is driven by E2F transcription factors released from hyperphosphorylated ppRB1 [22]. Hypophosphorylated pRB:E2F partially synthesizes CCNE. Like its predecessor, CCNE:CDK2 is reversibly regulated by WEE1 inhibition and CDC25A activation. CCNE degradation is controlled by SCF, which prevents re-entry into G1 [61]. The combined regulatory inputs create a bistable switch that commits the cell to DNA synthesis once CCNE activity exceeds a critical threshold. Then CCNA:CDK2 (**Fig. 1D**) becomes dominant during S phase, sustaining DNA replication and coordinating mitotic preparation. CCNA synthesis is induced downstream of E2F activation and repressed by APC/C:CDH1 until the G1/S transition is complete [21, 62]. Again, activity is regulated by WEE1 and CDC25A, with APC/CP:CDC20 ensuring its decline at the G2/M boundary [63, 64]. Finally, CCNB:CDK1 (**Fig. 1E**) governs mitotic entry and progression [51, 53]. CCNB accumulates steadily through G2, binding to CDK1. However, WEE1 kinase maintains inhibitory phosphorylation of CDK1 until it is dephosphorylated by CDC25C. Mitotic exit is driven by the sequential activities of the APC/C complex. First, APC/CP:CDC20 marks CCNB for degradation, initiating exit from mitosis. Then, APC/C:CDH1 becomes active in G0/G1 to clear residual CCNB and CDC20, thereby fully resetting the cycle.

**Figure 1.**
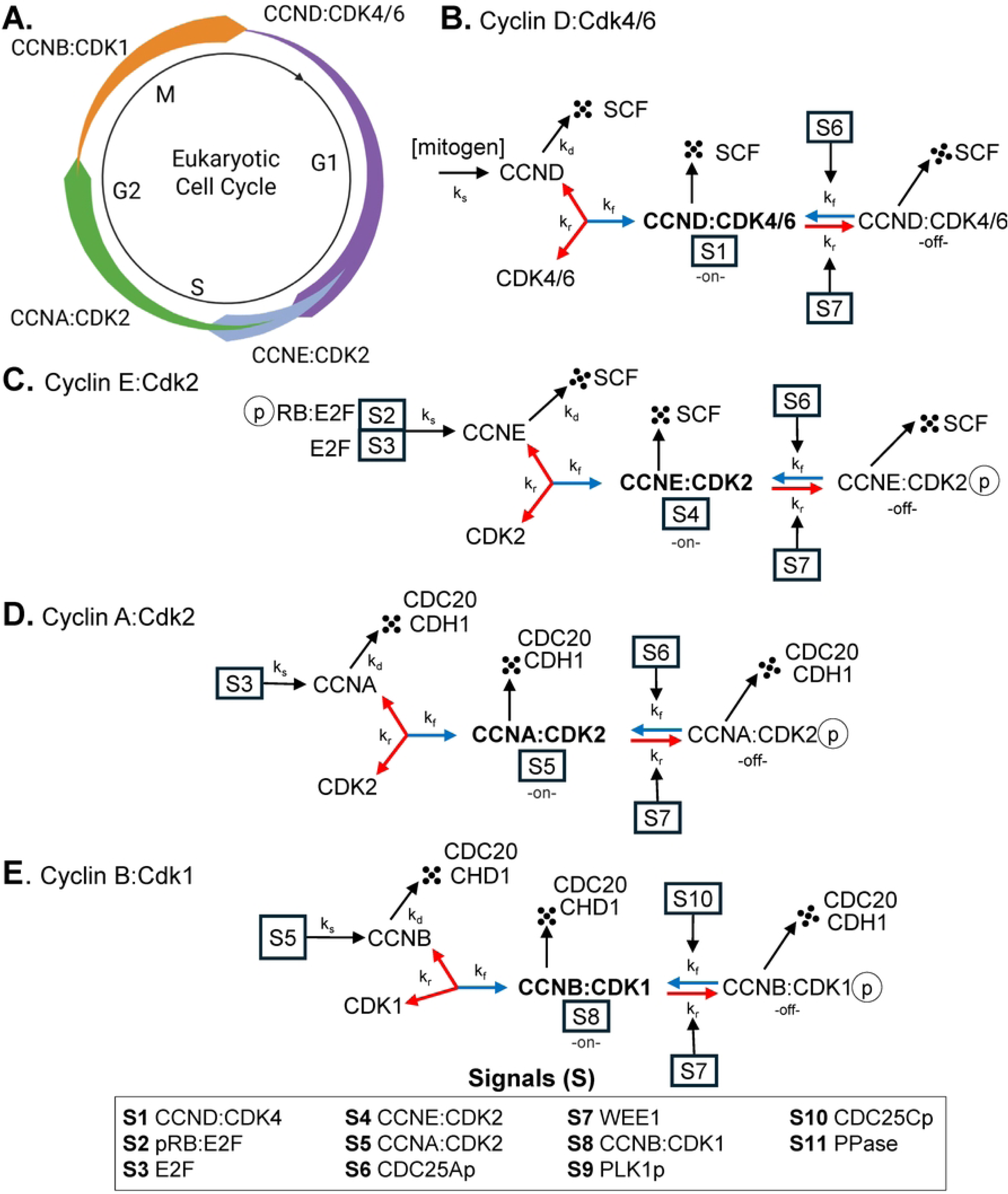
Schematic of the core cyclin-CDK network governing the eukaryotic cell cycle. The biochemical reaction network of key regulatory proteins is modeled as a system of ordinary differential equations (ODEs) based on mass conservation, using a hybrid framework that combines mass-action and Michaelis–Menten kinetics for reaction rates. The cycle is divided into four core regulatory modules corresponding to G1, S, G2, and M phases, each organized around a dominant cyclin–CDK complex that drives progression through its respective phase. Positive regulatory interactions are shown as blue arrows, inhibitory interactions as red arrows, and protein synthesis, degradation (terminating in a box for degradation), and (de)phosphorylation events. **(A)** Circular cartoon representation of the cell cycle phases showing the dominant cyclin–CDK complex active in each phase, with relative concentration profiles indicated. **(B)** G1 phase entry is driven by Cyclin D:CDK4/6, with Cyclin D synthesis stimulated by mitogenic signaling and degradation mediated by SCF ubiquitin ligase pathways. For ease of understanding, enzyme activities are noted as “on” or “off” and specific functional signals are indicated as S1-S11. **(C)** G1/S transition controlled by Cyclin E:CDK2, activated through CDC25A-mediated dephosphorylation and downregulated by SCF-dependent degradation to ensure unidirectional progression. **(D)** S-phase progression is maintained by Cyclin A:CDK2, which supports DNA replication and initiates G2 preparations before being targeted for degradation via APC/C:CDH1 and APC/CP:CDC20 to permit mitotic entry. **(E)** G2/M transition triggered by Cyclin B:CDK1. Inactive CCNB:CDK1p (also referred to as pre-MPF) is maintained by WEE1 kinase and rapidly activated by CDC25C phosphatase to initiate mitosis, followed by degradation via APC/C:CDH1 and APC/CP:CDC20 to allow mitotic exit. The model comprises coupled reaction modules that capture synthesis, degradation, and regulatory modification steps for each component, allowing simulation of coordinated oscillations and switch-like transitions across the cell cycle.

The activities of the Cyclin:CDK complexes are coordinated by several layers of regulation. We have limited regulations to changes in phosphorylation in this model (**Fig. 2A**). The WEE1 kinase phosphorylates CDKs, disrupting CDK binding to substrate. In turn, WEE1 activity is regulated, being suppressed by kinases (CCNA:CDK2, PLK1, CCNB:CDK1) and activated by phosphatases. Due to the involvement of numerous cellular phosphatase complexes, we have generalized these factors as “PPases” unless otherwise noted. Phosphorylated WEE1 is targeted for degradation by the SCF complex, ensuring its timely inactivation. In contrast to WEE1 activities, CDC25 phosphatases remove phosphates from CDKs. Together, WEE1 and CDC25 function as key regulatory nodes within feedback loops that drive phase transitions. Our model accounts for two isoforms of CDC25 (**Fig. 2A**). CDC25A synthesis is stimulated by E2F and its phosphatase activity is stimulated by CCNE:CDK2 and CCNA:CDK2 [65]. This mechanism creates a positive feedback loop that accelerates S-phase entry. The CDC25A phosphatase is marked for degradation by both APC/C:CDH1 and SCF, which ensures that this activation is transient. The CDC25C phosphatase operates similarly during G2/M, when activation by PLK1 and CCNB:CDK1 establishes another reinforcing loop. At the S/G2 boundary, PLK1 kinase is activated by CCNA:CDK2 and CCNB:CDK1, enabling it to stimulate CDC25C and inhibit WEE1, driving an irreversible commitment to mitosis [66–68]. At mitotic exit, PLK1 is targeted for degradation by APC/C:CDH1, ensuring timely inactivation of mitotic signaling [69]. Finally, general phosphatases (PPases) counteract CDK activity to reset the system. Together, these kinase and phosphatase modules create self-reinforcing feedback loops that sharpen phase transitions and preserve cell cycle fidelity.

**Figure 2.**
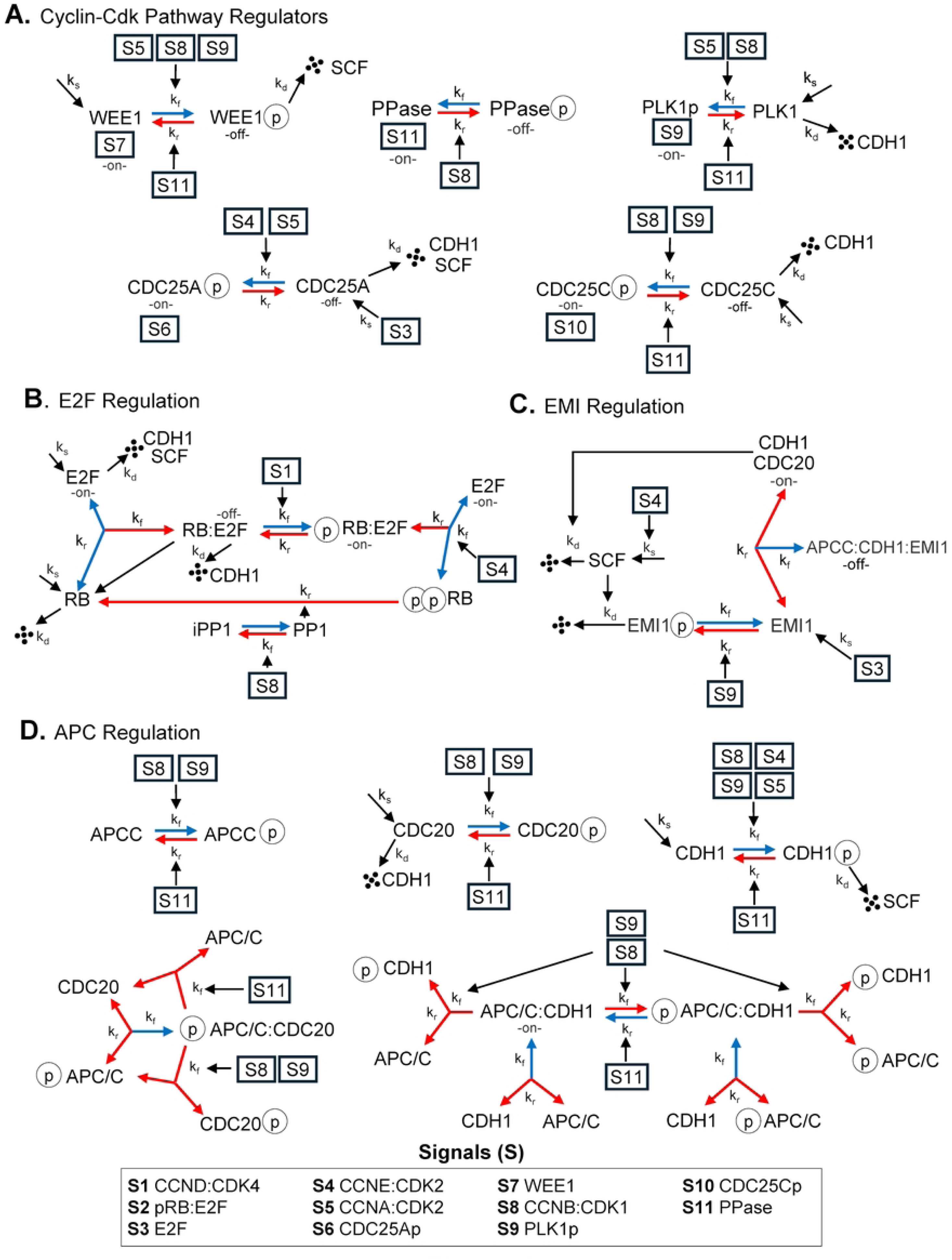
Detailed schematic of the regulatory signals controlling cell cycle progression. The signaling network governing cell cycle transitions with positive regulatory interactions are shown as blue arrows, inhibitory interactions as red arrows, and protein synthesis, degradation, and (de)phosphorylation events. For ease of understanding, enzyme activities are noted as “on” or “off” and specific functional signals are indicated as S1-S11. **(A)** Regulators of cyclin–CDK activity, including WEE1 kinase, generic protein phosphatases (PPase), PLK1 kinase, and CDC25A and CDC25C phosphatases. These factors control the activation state of CDK complexes by coordinated phosphorylation and dephosphorylation events, establishing precise timing for cell cycle transitions. **(B)** E2F regulatory network controlling transcription of S-phase–promoting genes. Positive and negative feedback loops integrate upstream CDK activity with transcriptional output to drive G1/S transition. **(C)** EMI1 and SCF regulation of APC/C:CDH1, including inhibitory interactions that prevent premature APC/C:CDH1 activation prior to mitotic exit. **(D)** APC/C regulatory pathways governing ubiquitin-mediated degradation of key substrates. Distinct branches activate APC/C(P):CDC20 during metaphase–anaphase transition and APC/C:CDH1 during mitotic exit, ensuring irreversible progression and cyclin degradation.

Progression into S phase is tightly controlled by the G1/S transition module, also referred to as the restriction point, which functions as a robust bistable switch centered on the RB1:E2F regulatory axis (**Fig. 2B**). In G1, the retinoblastoma protein (RB1) sequesters the E2F transcription factor, preventing the expression of S phase genes. Progression is initiated when active CCND:CDK4/6 begins to mono-phosphorylate RB1 (hypophosphorylated), which primes it for subsequent hyperphosphorylation by the CCNE:CDK2 complex [70]. This hyperphosphorylation (ppRB) breaks the RB1:E2F complex, releasing active E2F. ppRB is converted back to RB by the PP1 phosphatase. This mechanism creates a powerful positive feedback loop with released E2F stimulating synthesis of its own activator and leading to a switch-like activation that triggers a wave of transcription for S-phase genes such as CCNA and the APC/C inhibitor EMI1 [71].

Proteolysis is an essential driver of cell cycle directionality. Two major ubiquitin ligase systems coordinate this process: the SCF and APC/C complexes. The SCF complex is active from late G1 through S phase, targeting phosphorylated substrates such as CCND, CCNE, and CDK inhibitors for proteasomal degradation. This action ensures the timely destruction of G1 regulators. In contrast, the APC/C complex is the master regulator of G1 and mitosis. Its activity is suppressed during S and G2 by the inhibitor EMI1 (**Fig. 2C**). EMI1 expression is induced by E2F, and it binds directly to the APC/C complexes. This inhibition prevents the degradation of CCNA and CCNB, allowing them to accumulate. Later in mitosis, EMI1 is degraded by SCF, restoring APC/C activity. The activity of APC/C is temporally controlled by an ordered hand-off between its two co-activators, CDC20 and CDH1 (**Fig. 2D**). During the metaphase-to-anaphase transition in mitosis, high CCNB:CDK1 and PLK1 activities phosphorylate APC/C proteins [72]. These events enable APC/C to bind CDC20. The active APC/C:CDC20 complex marks Securin (PTTG1) and Cyclin B for degradation, initiating mitotic exit. CDH1 is held in an inactive (phosphorylated) state by CCNE:CDK2, CCNA:CDK2, PLK1p, and CCNB:CDK1. The resulting decline in CCNB:CDK1 activity allows for phosphatase-mediated dephosphorylation of CDH1 and formation of APC/C:CDH1 [73, 74]. This E3 ubiquitin ligase complex is active at the end of mitosis and throughout G1. Active APC/C:CDH1 marks S and M phase cyclins as well as CDC20 for proteasomal degradation [30, 75]. Together, these coordinated mechanisms establish an ordered sequence of synthesis, activation, inhibition, and degradation that underlies the oscillatory nature of the mammalian cell cycle.

### Initial model parameterization and stepwise refinement of the cell cycle dynamics

Initial model parameterization began with our previously validated mitotic module [50], providing a baseline set of 105 parameters governing core mitotic events. This framework was sequentially extended to incorporate regulatory mechanisms controlling G1, S, and G2 progression, including Cyclin:CDK-, E2F-, SCF-, and EMI-mediated pathways (**Appendix S2**). Regulatory modules were added incrementally to preserve mechanistic interpretability and to enable systematic evaluation of their impact on global cell cycle dynamics. After the addition of each regulatory layer, model behavior was assessed through a series of quantitative biological checks. To systematically explore feasible parameter regimes, we employed a constraint-based screening strategy in which parameter values were varied within predefined ranges around baseline estimates. For each parameter, ranges were specified through fixed percentage variations or explicit bounds, and the Cartesian product of these ranges was employed to construct the complete parameter grid. The resulting parameter grid was evaluated against multiple biological constraints, including correct temporal ordering of molecular events, maintenance of biologically realistic phase durations, and comparable activity scales across cyclin–CDK complexes. To further elaborate, we first verified the correct sequential activation of cyclin–CDK complexes, ensuring the ordered progression of protein complexes from CCND:CDK4 through CCNB:CDK1. Next, we evaluated the model’s ability to sustain autonomous oscillations by simulating extended time courses exceeding 10,000 cycles following a brief stabilization period. Oscillatory stability was quantified by measuring both the oscillation period and the peak CCNB:CDK1 activity, requiring less than 0.1% variation across cycles. Subsequently, we validated the timing of new mechanisms against biological literature. For example, APC/C:CDH1 activity was required to be active at the end of mitosis and persist through G1 [26]. Finally, we ensured that relative protein concentrations remained biologically plausible and that no single molecule’s concentration disproportionately dominate the system. Parameter sets that disrupted these timing relationships or produced disproportionate protein abundances were excluded from solution sets. This iterative refinement ensured that newly introduced mechanisms integrated coherently with existing dynamics rather than destabilizing the system.

The simulations are shown at a stable oscillatory, beginning at t = 4490, and showcase fully stabilized oscillations (**Fig. 3**). Vertical boundaries in each panel denote the phase transitions defined by molecular markers (**Fig. 3A**). Specifically, the border between G1 and S (G1/S) is characterized by peak Cyclin E levels, between S and G2 (S/G) is half-maximal PLK1 phosphorylation, between G2 and M (G2/M) is peak phosphorylated CDC25C, and between M/G1 is the rapid decline in phosphorylated Lamin A/C (LMNAp) marking mitotic exit. These boundaries closely correspond to experimental definitions of phase duration and order [67, 76–78]. The oscillatory dynamics of the four principal Cyclin:CDK complexes reveal a coordinated sequence of activation and decay (**Fig. 3A**). CCND:CDK4/6 initiates G1 entry and is gradually replaced by CCNE:CDK2 as the cell approaches the restriction point. As the S phase begins, CCNA:CDK2 becomes the dominant complex, sustaining DNA synthesis and promoting G1 progression. Finally, CCNB:CDK1 rises sharply at the G2/M transition to trigger mitotic entry, then rapidly collapses upon mitotic exit. The dynamics of free and complex Cyclin:CDKs show similar temporal trends across the cycle (**Fig. 3B–D**). Each complex displays a characteristic rise in Cyclin concentration (blue), followed by complex formation (green), while a pool of free CDK (red) remains present throughout, reflecting the balance between synthesis, degradation, and binding dynamics. The G2/M regulatory network (**Fig. 3E**) exhibits behavior comparable to that of the G2/M network but includes the activating and inhibitory control of CCNB:CDK1 by CDC25C and WEE1. During G2, inactive CCNB:CDK1 accumulates and is rapidly converted to its active form at the G2/M boundary. Activation of CDC25C and suppression of WEE1 establish a positive feedback loop that drives irreversible commitment to mitosis through the formation of active CCNB:CDK1. APC/C-mediated degradation of CCNB promotes mitotic exit, completing the oscillatory cycle. Together, these simulations demonstrate that the integrated regulatory network produces stable, self-sustaining oscillations in Cyclin:CDK activity that recapitulate the ordered progression of cell cycle phases observed experimentally.

**Figure 3.**
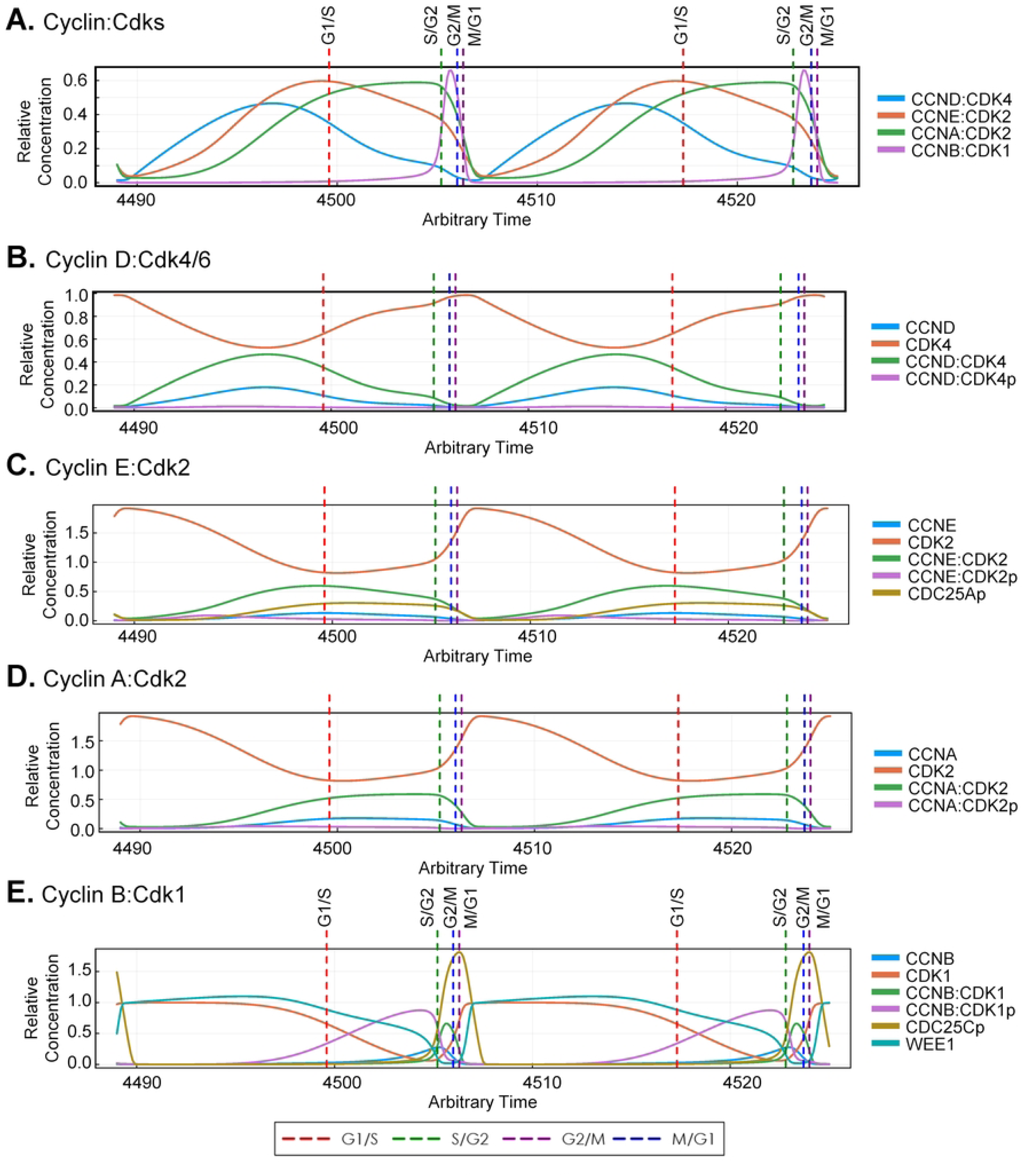
Dynamic simulation of core protein oscillations driving the eukaryotic cell cycle. Simulations of relative concentrations of key cell cycle proteins and their complexes are shown over a 40-hr timescale, encompassing two full cycles. The results are generated from the integrated ordinary differential equation (ODE) model, with each colored line representing a specific molecular species as defined in the panel legends. Vertical dashed lines indicate the major cell cycle transitions: G1/S (red), S/G2 (green), G2/M (purple), and M/G1 (blue). The biochemical reactions, ODEs, and optimized model parameters are presented in the Supporting Information. **(A)** Sequential oscillations of the four core cyclin–CDK complexes (Cyclin D:CDK4/6, Cyclin E:CDK2, Cyclin A:CDK2, Cyclin B:CDK1) over two cycles. **(B)** Accumulation of Cyclin D:CDK4/6 activity (green line) in early G1 promotes progression toward the restriction point and primes Cyclin E expression. **(C)** Peak Cyclin E:CDK2 activity (green line) at the G1/S transition initiates DNA replication, followed by rapid Cyclin E degradation to ensure unidirectional progression. **(D)** Cyclin A:CDK2 activity (green line) rises through S-phase to maintain replication and initiate early G2 functions, declining before mitosis. **(E)** Cyclin B:CDK1 accumulates in G2 as inactive CCNB:CDK1p (purple line), maintained by WEE1 kinase, and is rapidly activated (green line) by CDC25C at G2/M to trigger mitotic entry.

We next examined the coordination between synthesis and degradation pathways. This includes activities of the three major E3 ubiquitin ligase complexes, SCF (green), APC/C:CDH1 (blue), and APC/CP:CDC20 (red) (**Fig. 4A**). The SCF complex is active primarily in S and G2 phases but reaches a maximum after mitosis. The SCF complex is then marked for degradation by APCC:CDH1 as the cell cycle reverts to its quiescence state. In contrast, APC/CP:CDC20 and APC/C:CDH1 act sequentially during mitosis and mitotic exit. APC/CP:CDC20 is activated by PLK1- and CCNB:CDK1-mediated phosphorylation. With the decline in CCNB levels, APC/C:CDH1 becomes the dominant ligase and clears residual mitotic substrates. Our model shows active APC/C:CDH1 throughout G1 with decreasing concentrations into the S phase. The regulation of EMI1 (**Fig. 4B**), an inhibitor of APC/C, plays a pivotal role in restricting APC/C:CDH1 activity at the G1/S transition. EMI1 accumulates downstream of E2F activation and binds to APC/C:CDH1, suppressing its activity as cells approach the G1/S boundary. As PLK1 phosphorylation increases during S and G2 phases, EMI1 becomes progressively inactivated, relieving APC/C inhibition. The model also captures a modest inhibitory effect of EMI1 on APC/CP:CDC20 during S phase, preventing premature accumulation of mitotic regulators prior to entry into mitosis. Synthesis of factors is driven, in part, by activation of the E2F transcription factor. (**Fig. 4C**). Free E2F activity rises rapidly during early G1, paralleling the accumulation of CCND and the decline of APC/C:CDH1. Levels peak near the G1/S transition and decline by the end of S phase as feedback inhibition is restored. The RB1:E2F complex undergoes progressive phosphorylation, resulting in low relative concentrations of bound RB1:E2F throughout most of the cycle. Total RB1 levels remain relatively stable, peaking at the end of mitosis, when dephosphorylation restores the inactive form of RB1.

**Figure 4.**
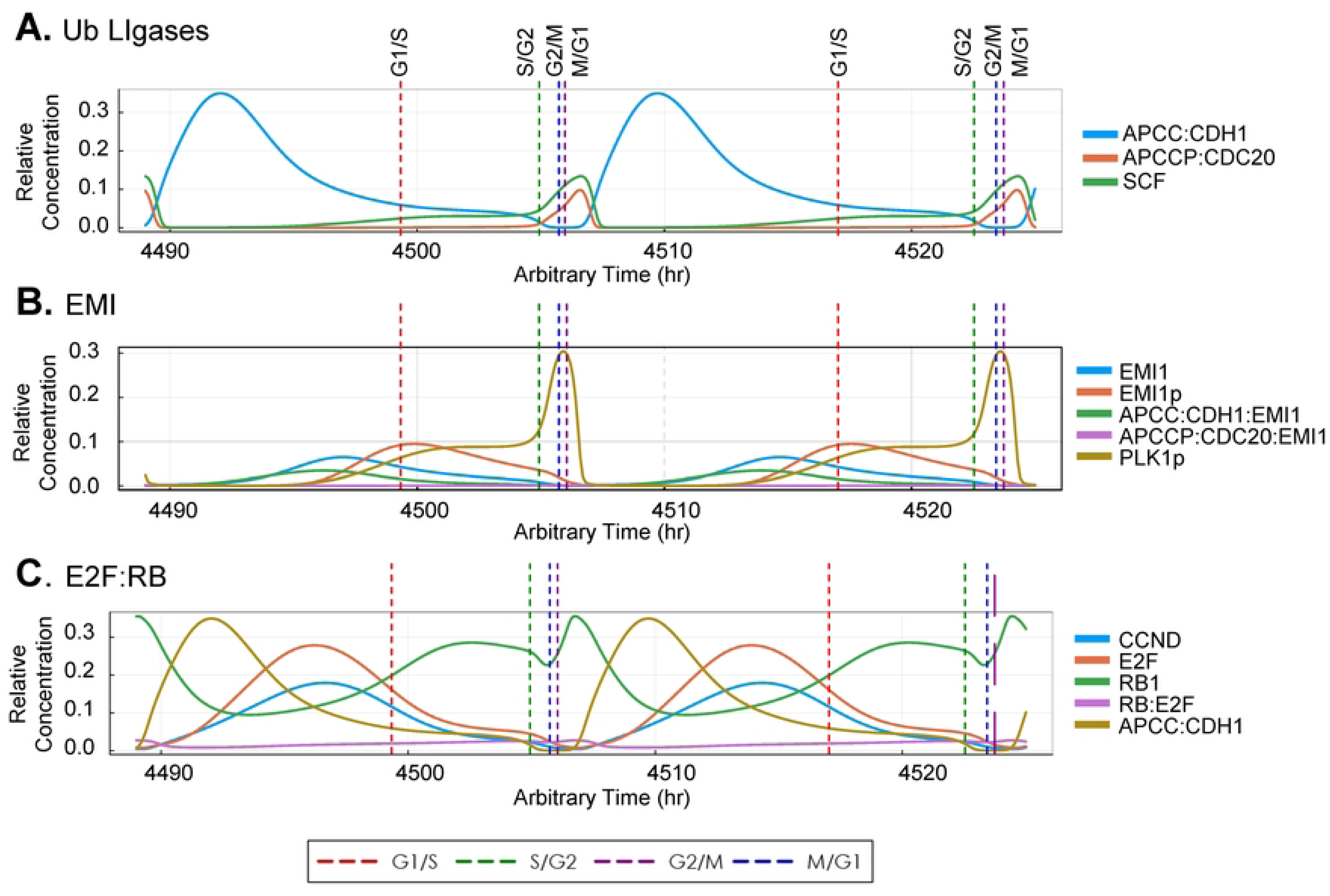
Simulated dynamics of core regulators across the full eukaryotic cell cycle. Relative concentrations for key components of three regulatory pathways are shown over a 40-hour timescale, encompassing two full cell cycles. The results are generated from the integrated ordinary differential equation (ODE) model, with each colored line representing a specific molecular species as defined in the panel legends. Vertical dashed lines indicate the major cell cycle transitions: G1/S (red), S/G2 (green), G2/M (purple), and M/G1 (blue). **(A)** Ubiquitin ligase network dynamics, highlighting the timed activation and degradation of APC/C and SCF complexes that coordinate substrate turnover during specific cell cycle phases. **(B)** EMI pathway regulation, showing the inhibition of APC/C:CDH1 by EMI1 and its relief by PLK1 at mitotic entry to permit substrate degradation. **(C)** E2F:RB pathway dynamics, illustrating RB-mediated repression of E2F transcription factor in early G1 and activation following phosphorylation by cyclin–CDK complexes promoting S-phase gene expression.

To evaluate the accuracy and robustness of the cell cycle model, we first tested its ability to reproduce canonical cell cycle checkpoint responses under perturbations. These checkpoints are critical control nodes that ensure fidelity of division and are frequently dysregulated in cancer and other diseases. We began by simulating the cell cycle’s dependence on extracellular mitogenic signals over multiple rounds of division (**Fig. 5A**). Mitogen withdrawal was modeled by turning off Cyclin D synthesis, thereby eliminating CCND:CDK4/6 activity. This disrupted CCND:CDK4 formation, stabilized hypophosphorylated RB1, and repressed E2F-driven transcription, leading to a G1-arrested state in which oscillations of downstream Cyclin:CDK complexes ceased entirely (**Fig. 5B, 5C**). This response closely mimics serum starvation, in which cells deprived of mitogenic growth factors exit the cell cycle and enter quiescence. Upon reintroduction of the mitogen signal after 50 hrs (**Fig. 5B**) and 100 hrs (**Fig. 5C**), the model immediately reinitiated Cyclin D synthesis, RB1 phosphorylation, and E2F activation, resulting in the rapid recovery of oscillations. The speed and fidelity of this recovery demonstrate that the model captures both the reversible nature of the restriction point and the bistable logic governing G1 entry and exit.

**Figure 5:**
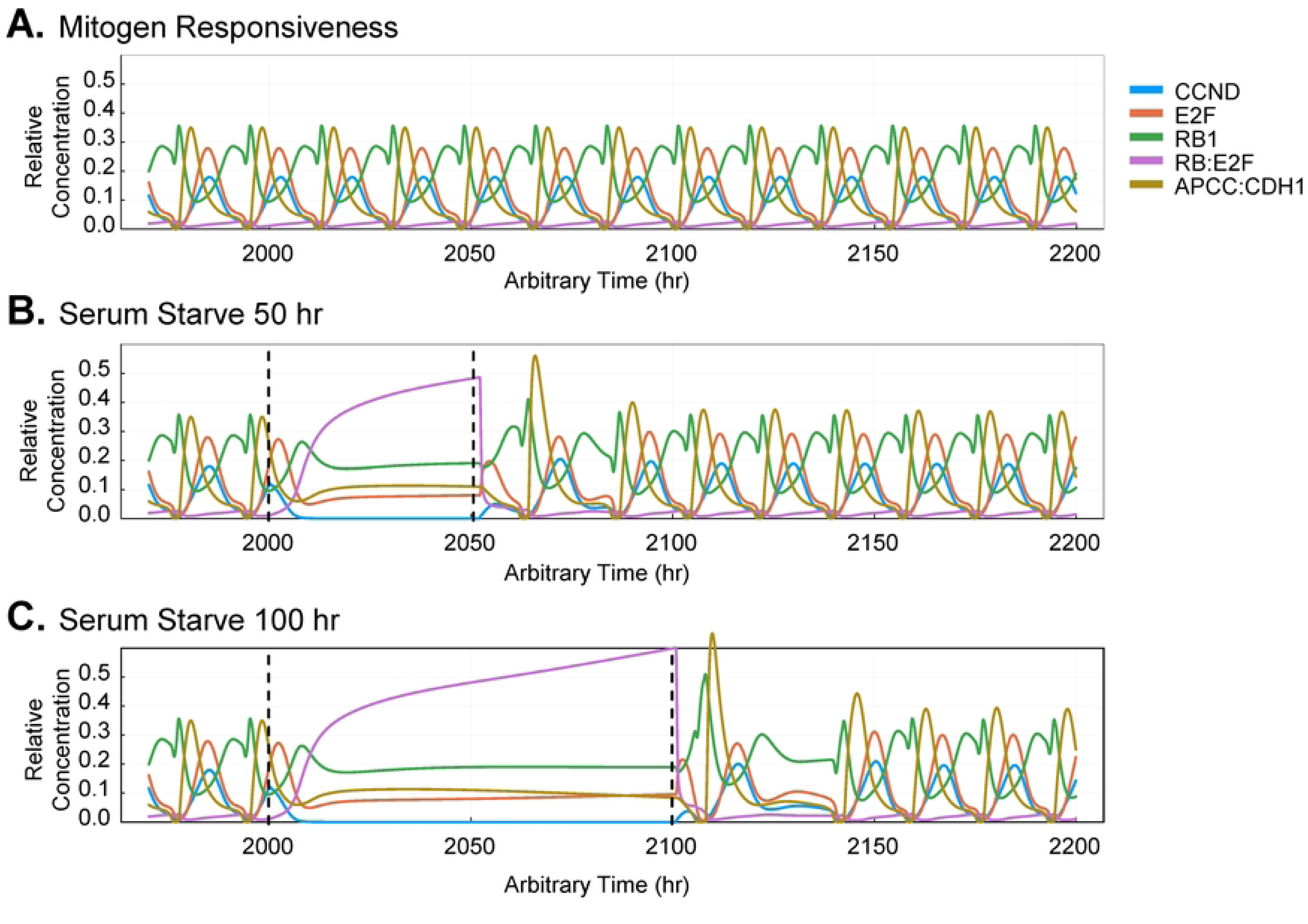
Simulated reversible G1 arrest and cycle re-entry in response to transient mitogen withdrawal. Simulations using the ODE model illustrate the cell cycle’s response to temporary removal and subsequent reintroduction of a mitogen growth signal, mimicking *in vitro* serum starvation. **(A)** Under constant mitogen stimulation, the system maintains continuous and stable oscillations, with sequential peaks of Cyclin D driving repeated divisions. Thirteen cycles are shown. **(B-C)** We simulated mitogen withdrawal by preventing cyclin D synthesis. Following withdrawal (first vertical dashed line), cyclin oscillations stop, which is consistent with an arrest for **(B)** 50 hr and **(C)** 100 hr. Restoration of the signal (second vertical dashed line) results in stable oscillations indistinguishable from the control.

Next, we simulated cellular responses upon the introduction of genotoxic stress. A DNA damage response (DDR) module was previously added (**Fig. 6A**) [51], where double-strand breaks trigger activation of Ataxia-Telangiectasia Mutated kinase (ATM). Activated ATM phosphorylates and activates Checkpoint Kinase 2 (CHK2), which in turn phosphorylates and stabilizes the tumor suppressor protein TP53 [79–81]. TP53 levels are further governed by a negative feedback loop with its E3 ligase, MDM2. MDM2 targets unphosphorylated TP53 for proteasomal degradation, which is transcriptionally induced by phosphorylated TP53 [82]. The primary downstream effector of this pathway is the cyclin-dependent kinase inhibitor p21 (CDKN1A), which is synthesized in a TP53-dependent manner [79]. Following its induction, p21 binds to and inhibits Cyclin:CDK complexes [83–86]. We simulated a DNA damage response by introducing a transient pulse of ATM activity to mimic a double-strand break event during CCNE:CDK1 activities (**Fig. 6B**). The model accurately propagated this perturbation through the downstream cascade with elevated p21 inhibiting multiple cyclin–CDK complexes, including CCNE:CDK2, CCNA:CDK2, and CCNB:CDK1 (**Fig. 6C**). Importantly, this checkpoint was reversible. Once ATM activity returned to baseline, p53 and p21 levels declined, CDK activity was restored, and the simulation resumed normal cycling (**Fig. 6B-C**). Together, these tests confirmed that the model’s regulatory architecture is robust and can reproduce critical checkpoint behaviors.

**Figure 6.**
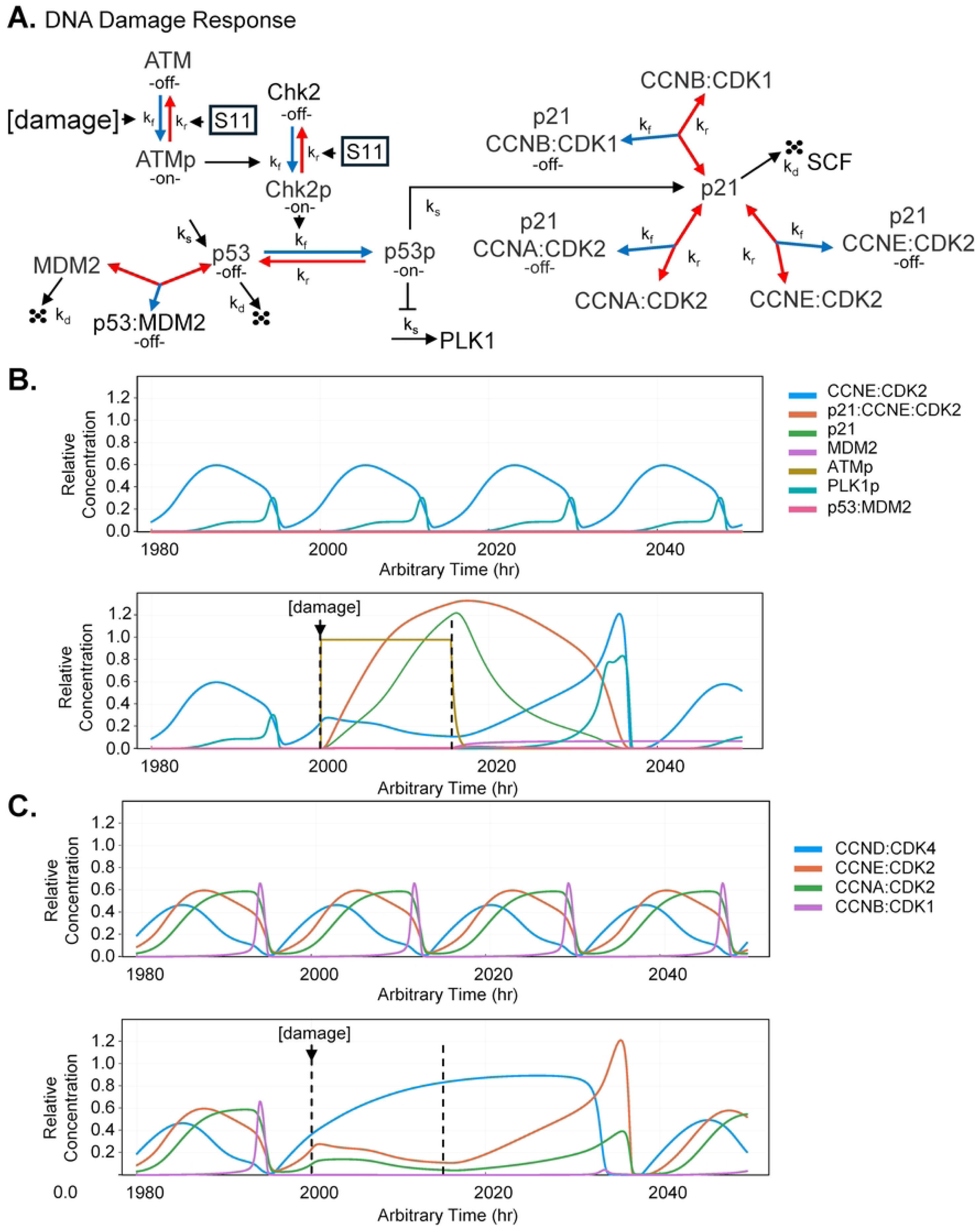
Simulation of DNA damage-induced cell cycle arrest and recovery. Predicted dynamics of key proteins in the double-strand break DNA damage response (DDR) pathway. **(A)** The modeled DDR network [51] is initiated by activation of ATM (ATMp), which phosphorylates and activates Chk2 (Chk2p). Activated Chk2p, in turn, phosphorylates and stabilizes the transcription factor p53 (p53p), which induces the expression of the CDK inhibitor p21. p21 binds and disrupts the activities of cyclin–CDK complexes. Within the pathway, p53 is negatively regulated by MDM2. For ease of understanding, activities are noted as “on” or “off.” Red arrows indicate inhibition, while blue arrows indicate activation. **(B)** *In silico* simulation of ATM activation (ATMp) (lower panel, dashed lines) resulting in increased p21 (green line) in complex with Cycle E:CDK2 (red line) compared to normal oscillations (upper panel). **(C)** Inactivation of ATM (second dashed line) restores normal cycle-CDK oscillations after a short period of time.

### Aligning and predicting cell cycle phases using human glioma cells

In addition to developing a stable, responsive, and oscillatory human cell cycle model, we aimed to ensure that the temporal dynamics were grounded in experimental data. We applied the live-cell FastFUCCI (fluorescence ubiquitination-based cell cycle indicator) reporter system, previously developed by Koh *et al.* [52], to track cell cycle phases (**Fig. 7**). The approach uses oscillations in CDT1 and Geminin fused to fluorescent proteins. We used the human U251-MG glioma cell line for these studies. We transduced cells with the reporter, selected for expression, and enriched for fluorescing cells by limiting dilution. Using live-cell imaging collected every 2 hrs over a 96 hr period, we tracked individual cells, where a specific cell gives rise to daughter cells (e.g., Cell #1 to #1.1 and #1.2, then #1.1 to #1.1.1 and #1.1.2, etc.) (**Fig. 7A**). The reporter system visually distinguishes single cell G1 (CDT1-red) from S/G2/M (Geminin-green) phases. Cells in mitosis exhibit high levels of Geminin (green) and are physically rounded, followed by two non-fluorescing daughter cells at the next imaging time point. CDT1 (red) and Geminin (green) protein levels are regulated by SCF and APC/C:CDH1 complexes, respectively, and we incorporated these signals into our model (**Fig. 7B**). CDT1 is a licensing factor required for DNA replication that accumulates during G1 and is degraded at the onset of S phase, while Geminin inhibits DNA replication licensing by blocking CDT1 and is expressed from S phase through mitosis [87]. We elected to set t=0 hrs after observing the first mitotic event giving rise to daughter cells (i.e., t=0 starting at Cell #1.1). Live-cell imaging and counting revealed a mean doubling time of approximately 18 hrs (**Fig. 7C)**, with mean G1 lasting 10 hrs and S/G2 lasting 8 hrs (**Fig. 7E**). Mitosis is a rapid event and occurs faster than we can capture using a 2 hrs imaging time points; therefore, we set M as 2 hrs for these studies.

**Figure 7.**
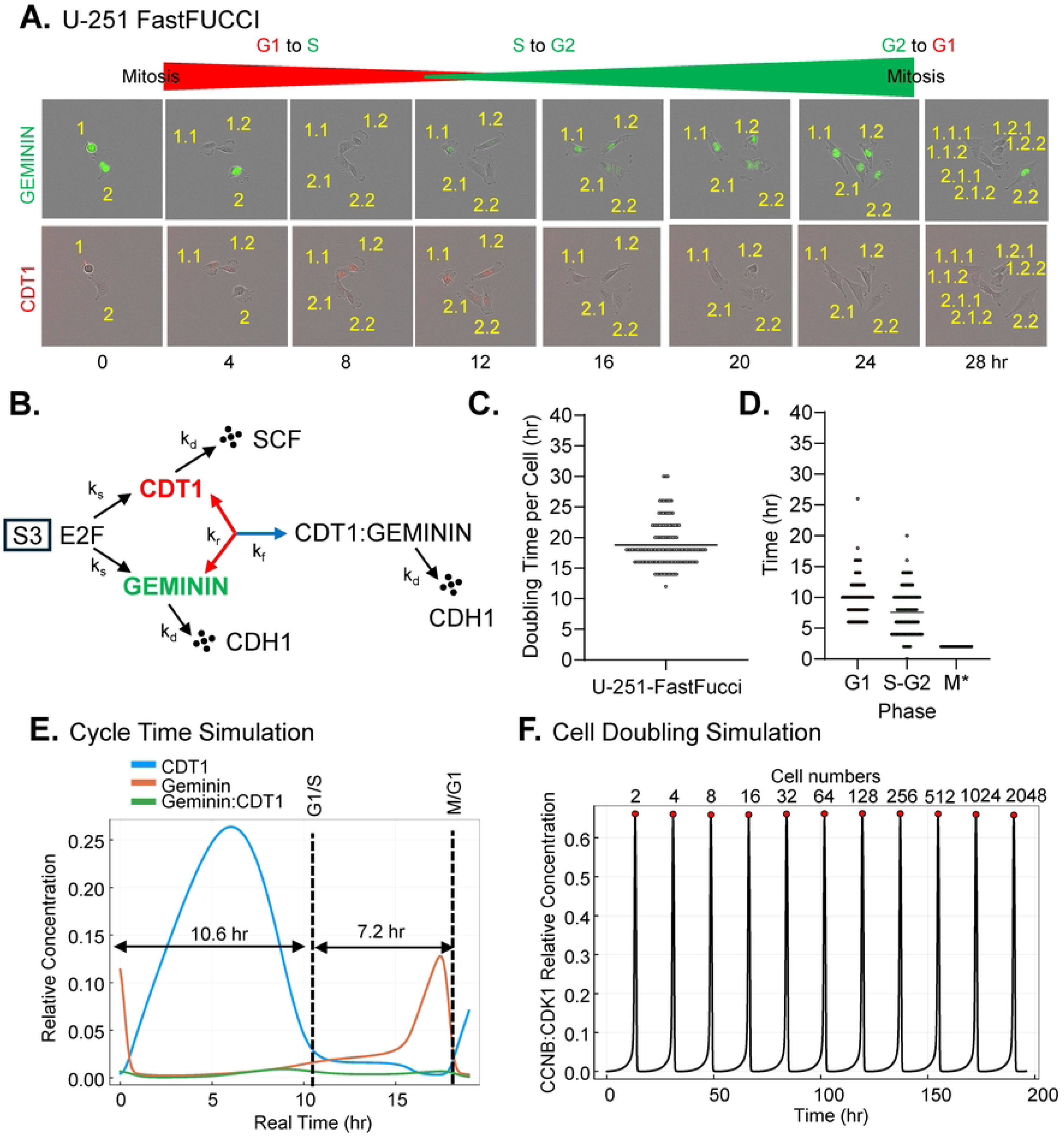
Experimental validation of the cell cycle model using FastFUCCI system and CDT1-Geminin simulation. **(A)** Representative time-lapse microscopy images from select time points of U251-MG cells expressing the FastFUCCI reporter [52]. Cell cycle phases are visualized by fluorescent protein fusions with CDT1 (red, marking G1 phase) and Geminin (green, marking S/G2/M phases). Individual cells are tracked to quantify phase durations (e.g., Cell #1 to #1.1 and #1.2, then Cell #1.1 to #1.1.1 and #1.1.2, etc.). **(B)** A schematic of the key regulatory interactions governing oscillations of CDT1 and Geminin. The network includes regulation of CDT1 and Geminin by E2F (S3 signal), SCF, and APC/C:CDH1. **(C)** Quantification of cell cycle doubling time from the FastFUCCI imaging data. Individual cells were tracked, and timing was determined following the first cell division. Each data point represents the transition time for a single cell (e.g., Cell #1 to Cell #1.1 and #1.2). These data show a mean doubling time of 18 hrs. (D) Quantification of cell cycle phase durations from the FastFUCCI imaging data with an average G1 time of 10.5 hrs. and average S/G2 7.5 hrs. Mitosis has been defined by cell rounding with high Geminin expression becoming two non-fluorescent daughter cells, which occurs in <2 hrs. **(E)** Model simulation showing relative concentrations of CDT1, Geminin, and the CDT1:Geminin complex over one complete cycle in hours. The model predicts a G1 phase duration (high CDT1) of 10.55 hrs and an S/G2/M phase duration (high Geminin) of 7.18 hrs, resulting in a doubling time of 17.73 hrs. **(F)** The model simulates a consistent doubling time over many generations, confirming the integrity of the simulated cell cycle.

We extracted phase transition times and G1/S and M/G1 boundaries from the experimental data and used them as target variables for model calibration. During parameter screening, candidate parameter sets were required to reproduce these phase transition times within a predefined tolerance window, allowing deviations of ±0.3 hrs around the experimentally observed values. The optimized model achieved a stable simulated doubling time of 17.73 hrs, with G1 and S/G2/M durations of 10.55 hrs and 7.18 hrs, respectively (**Fig. 7E**). The model maintained this precise timing over many generations, confirming the stability and accuracy of the calibrated oscillator. We calculated doubling time, defined as the interval between successive mitoses in the same tracked cell (first to second division). These timings were averaged among all tracked cells. In the simulation, doubling time is calculated by detecting peaks in the CCNB:CDK1 concentration signal (**Fig. 7F**). The algorithm identifies local maxima using peak-detection parameters (height ≥ 0.2, prominence ≥ 0.2) to filter out noise. The time intervals between consecutive peaks represent doubling time, with each peak contributing to the total cell numbers. In summary, we are able to calibrate our model to the average doubling time and phase times for a given cell type using data from *in vitro* studies.

### Predicting drug-mediated inhibition of cell proliferation

We experimentally tested and simulated the effects of two cell cycle inhibitors, abemaciclib and volasertib, using our validated cell cycle model and then compared the results. Abemaciclib is a CDK4/6 inhibitor that blocks phosphorylation of RB, preventing the release of the E2F transcription factor and arresting cells at the G1/S checkpoint [88–90]. We treated FastFUCCI-labelled U251-MG glioblastoma cells with abemaciclib at 0.01, 0.1, or 1.0 μM, including a DMSO control. We analyzed the impact on cell proliferation by counting cells at specific time points (**Fig. 8A**) and by visualizing cells using live cell imaging (**Fig. 8B**). We observed that increasing concentrations of abemaciclib resulted in reduced cell numbers over time (**Fig. 8A**) which is reflected in the calculated doubling times (**Fig. 8C**). In comparison to DMSO at 23 hrs, average doubling time with 0.01 µM abemaciclib was 25 hrs and with 0.1 µM was 42 hrs (**Fig. 8C**). Addition of 1.0 µM abemaciclib resulted in an arrest and no cell doublings (**Fig. 8A,C**). We also analyzed drug impact using live cell imaging, which showed increasing concentrations resulted in fewer cells over time, including accumulation of vacuoles in the cytoplasm at 1.0 µM (**Fig. 8B**). When averaging doubling times per cell (**Fig. 8D**), we observed an average of 20 hrs for DMSO, 26 hrs for 0.01 µM abemaciclib, 28 hrs for 0.1 µM, and no cell division at 1.0 µM. Using FastFUCCI, we observed increasing numbers of cells accumulating in G1 upon increasing abemaciclib concentrations, starting at 10 hrs for DMSO to 15 hrs and 21 hrs at 0.01 µM and 0.1 µM, respectively (**Fig. 8E**). In contrast, the averaged time for S-G2-M was 12 hrs for all conditions during proliferation (**Fig. 8F**). Since no doubling was observed at 1.0 µM abemaciclib, we wanted to track what happens to individuals cells (**Fig. 8G**). Cells starting in G1 fluctuated into S/G2 (Geminin) and back to G1, while cells starting in S/G2 could divide but subsequently arrest in G1. On a rare occasion, a cell was observed to progress during drug exposure (**Fig. 8G**), suggesting the possibility of resistance. Overall, inhibition of cell division is concentration-dependent and consistent with the known activity of abemaciclib, predominately inducing an arrest in G1.

**Figure 8.**
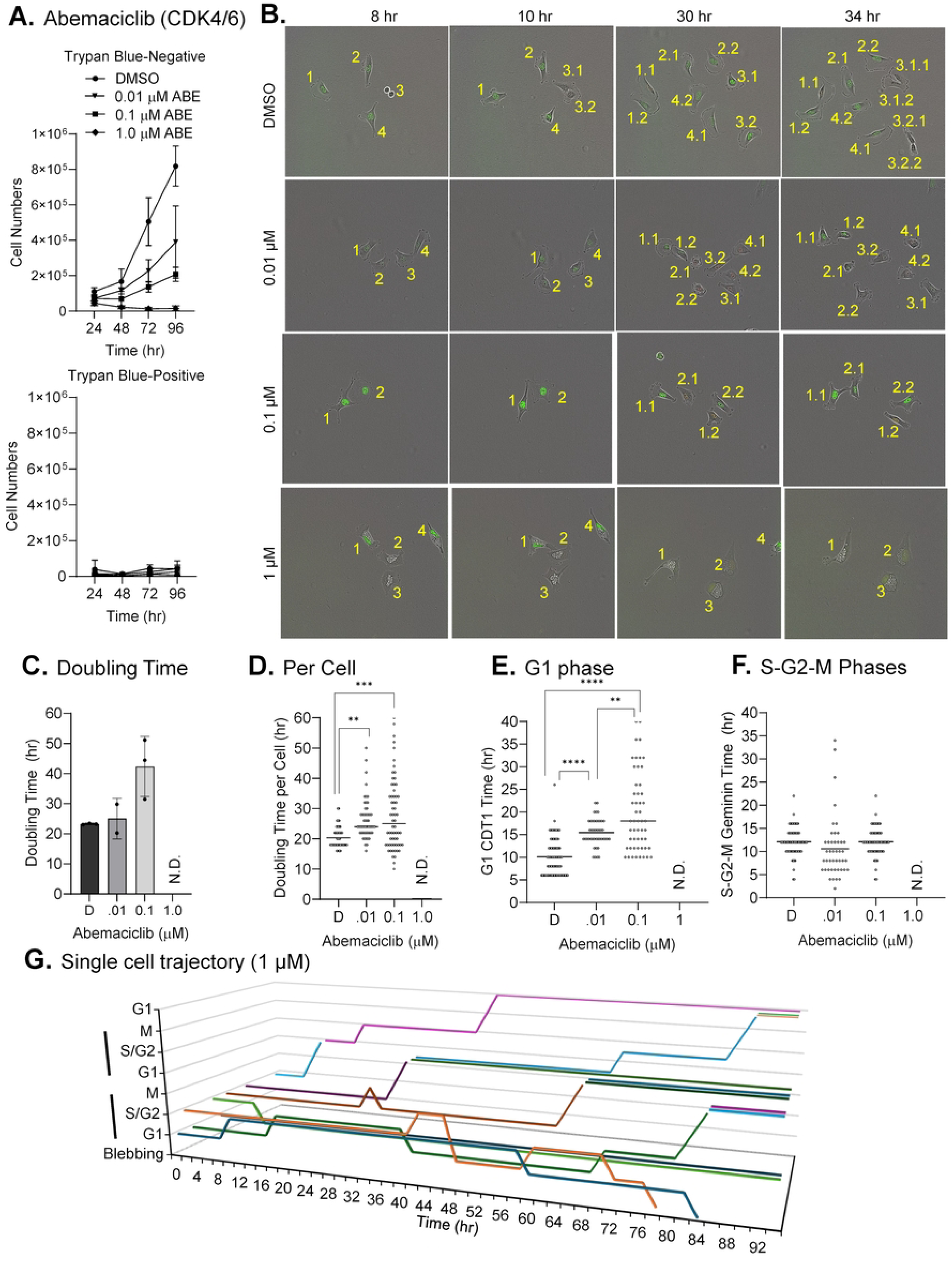
Dose-dependent arrest following CDK4/6 inhibition by abemaciclib. **(A)** U251-MG FastFUCCI cells were plated subconfluently and treated with the indicated concentrations of abemaciclib or DMSO. Media with inhibitor was replaced every 24 hrs. Cell numbers were determined using a hemocytometer and trypan blue staining. Trypan-blue negative (top panel) and positive (bottom panel) cell numbers are shown (n=3). **(B)** Representative time-lapse live-cell imaging of U251-MG FastFUCCI cells treated with vehicle control (DMSO) or increasing concentrations of abemaciclib. Individual cells and their daughter cells are numerically identified. **(C)** Calculated cell doubling times using data from (A) with no doubling time determined 1.0 µM abemaciclib (N.D., not determined). **(D)** Doubling times for individual cells across treatments. **(E)** Phase durations of single cells in G1 (CDT red) and (F) S/G1/M (Geminin green) across treatments. **(G)** Graph showing representative cell cycle dynamics for cells treated with 1.0 µM abemaciclib. Each line represents phase changes for a single cell over time and any resulting daughter cells. Individual cells were tracked by cell cycle phase color. Cell blebbing, consistent with apoptosis, is noted. n=3 biological replicate experiments with standard deviation from the mean. Standard deviation from the mean is shown, and significance is determined by Student’s t-test.

Computationally, abemaciclib was modeled as an inhibitor of RB:E2F phosphorylation using a Hill-type equation (Appendix 3, Eq. 3.0.1), slowing E2F accumulation and lengthening the cell cycle. Parameters for Hill-type equation were calibrated using experimental doubling times from live cell imaging analysis, with percent change relative to controls as the fitting variable to account for baseline variability. Simulations predicted doubling times of 17.7 hrs for DMSO, 17.7 hrs at 0.01 μM abemaciclib, and 23.9 hrs at 0.1 μM (**Fig. 9A-C**). At 1.0 μM, the model predicted a stalled G1 state with cessation of oscillations which matches experimental outcomes. Finally, we compare the cell doubling times between our different analyses (**Fig. 9D**). This demonstrates increasing doubling times for 0.1 μM abemaciclib with an arrest at 1.0 μM for all analyses including the simulated responses (**Fig. 9D**).

**Figure 9.**
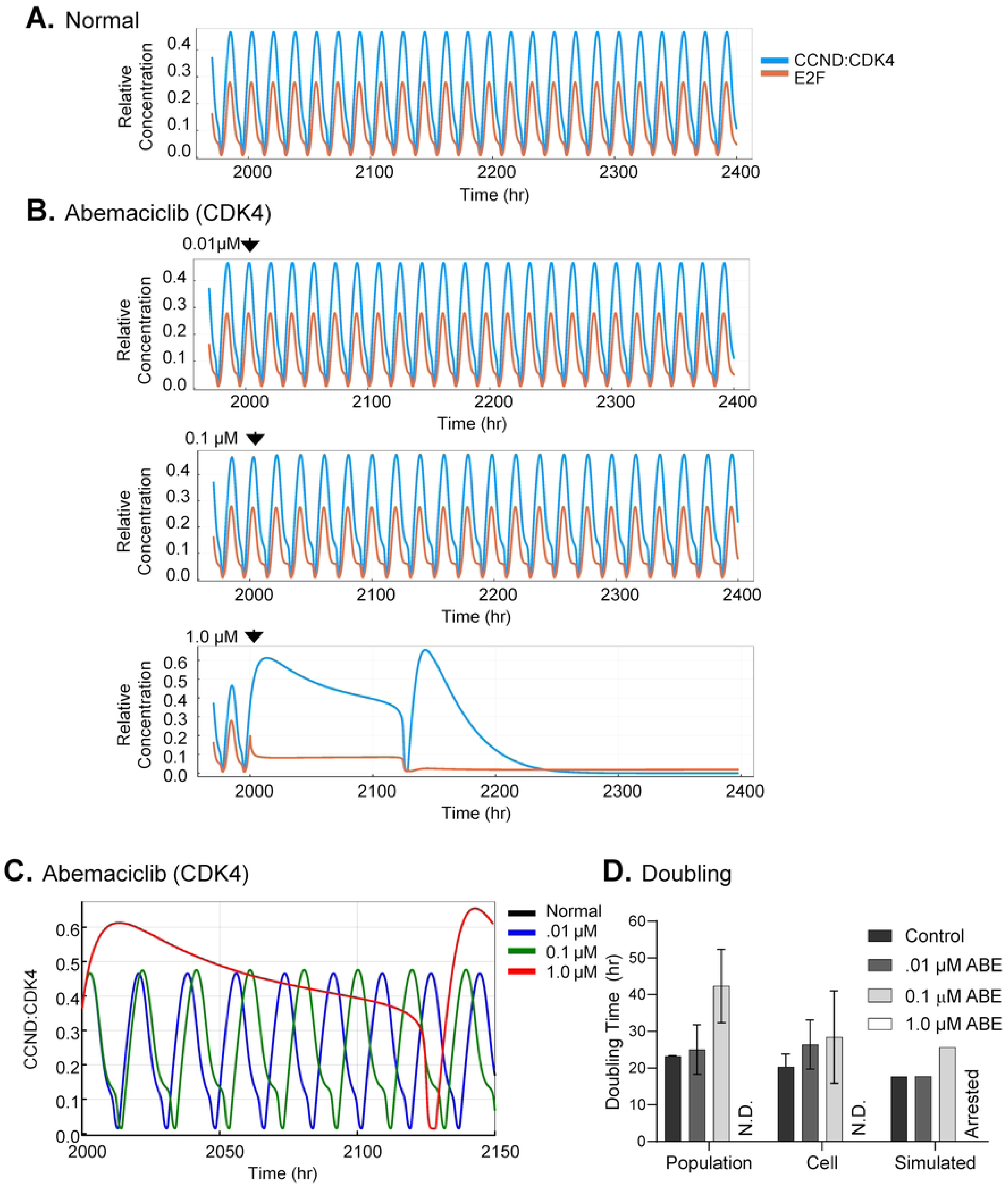
Model responsiveness to abemaciclib-mediated inhibition of CCND:CDK4 in silico. **(A)** Baseline simulation of G1 phase regulators, CCND:CDK4 and E2F, under normal conditions, depicting stable, periodic oscillations over time. **(B)** Simulated response to abemaciclib treatment at increasing concentrations (0.01 µM, 0.1 µM, 1.0 µM), demonstrating a dose-dependent slowing of the cell cycle oscillations. As experimentally observed, simulation with 1.0 µM abemaciclib results in loss of oscillations. **(C)** Overlay of CCND:CDK4 oscillations from control and treated simulations in (A) and (B). **(D)** Quantitative validation comparing doubling times determined by *in vitro* counting cell numbers of the population, *in vitro* single cell dynamics, and *in silico* simulations under all treatments.

Next, we analyzed the effect of the PLK1 inhibitor, volasertib, on cell proliferation. Inhibition of PLK1 disrupts spindle assembly and centrosome maturation, causing G2/M arrest and mitotic catastrophe [91]. We treated the FastFUCCI-labelled U251-MG glioblastoma cells with DMSO control or volasertib at 1.0 nM, 2.5 nM, or 5.0 nM, and again analyzed its impact by counting cells at specific time points (**Fig. 10A**) and live cell imaging (**Fig. 10B**). Increasing concentrations of volasertib resulted in reducing cell numbers over time compared to vehicle control (**Fig. 10A**). The calculated cell doubling times using these data showing 21 hrs for DMSO, 26 hrs for 1.0 nM volasertib, 28 hrs for 2.5 nM, and no cell doubling at 5.0 nM (**Fig. 10C**). Similar reductions in cell doubling were evident using live cell imaging with 5.0 nM volasertib resulting in Geminin-positive rounded cells (**Fig. 10B**, yellow numbering) as well as cell blebbing, all indicators of mitotic catastrophe (**Fig. 10B**, red numbering). By tracking individual cells, we calculate a mean doubling time of 21 hrs for DMSO, 26 hrs for 1.0 nM, and 28 hrs for 2.5 nM volasertib (**Fig. 10D**). We quantified disruptions in both cell cycle phases for treated cultures with increasing G1 and S-G2-M (**Fig. 10E**) with increasing time expressing Geminin with higher volasertib (**Fig. 10F**). We subsequently analyzed what happens to cells at 5 nM volasertib (**Fig. 10G**). This shows that most cells enter M phase and subsequently undergo blebbing, again an indicator of mitotic catastrophe. Overall, these data demonstrate increased cell doubling times following disruptions across all cell cycle phases, with the highest volasertib concentrations resulting in mitotic catastrophe.

**Figure 10.**
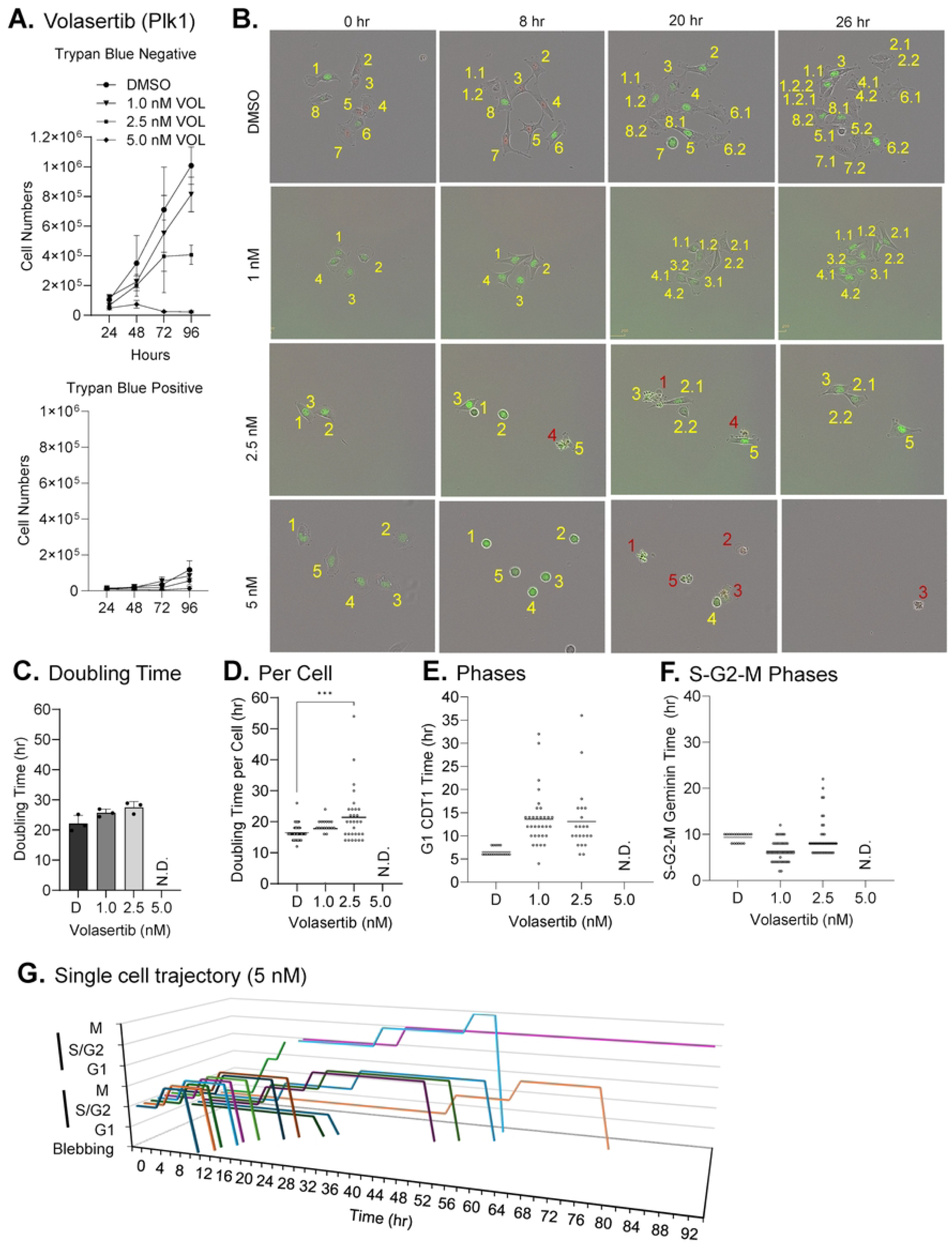
Inhibition of PLK1 by volasertib results in mitotic arrest. **(A)** U251-MG FastFUCCI cells were plated subconfluently and treated with the indicated concentrations of volasertib or DMSO. Media with inhibitor was replaced every 24 hrs. Cell numbers were determined using a hemocytometer and trypan blue staining. Trypan-blue negative (top panel) and positive (bottom panel) cell numbers are shown. n=3 **(B)** Representative time-lapse live-cell imaging of U251-MG FastFUCCI cells treated with vehicle control (DMSO) or increasing concentrations of volasertib. Individual cells and their daughter cells are numerically identified (yellow) with cells exhibiting signs of cell death indicated (red) **(C)** Calculated cell doubling times using data from (A) with no doubling time determined at 5.0 nM volasertib (N.D., not determined). n=3 **(D)** Doubling times for individual cells across treatments. **(E)** Phase durations of single cells in G1 (CDT red) and S/G1/M (Geminin green) across treatments. **(F)** Graph showing representative cell cycle dynamics in cells treated with 5.0 nM volasertib. Each line represents phase changes for a single cell over time. Cell blebbing was observed, suggestive of mitotic catastrophe and cell death. Standard deviation from the mean is shown, and significance is determined by Student’s t-test.

Computationally, volasertib was modeled by inhibiting PLK1 phosphorylation of CDC20, CDH1, APC/C, WEE1, and CDC25C using a Hill-type equation (Appendix 3, Eq. 3.0.2), delaying mitotic progression and elevating Cyclin B:CDK1 levels. This is more complex than abemaciclib, which disrupts only CDK4/6 phosphorylation of RB. Inhibition parameters were calibrated using experimental data with percent change in doubling time as the fitting variable. Simulations predicted doubling times of 17.7 hrs for DMSO (**Fig. 11A**) and 17.7 hr and 23.6 hrs at 1.0 nM and 2.5 nM volasertib, respectively (**Fig. 11B,C**). At 5.0 nM volasertib, the model predicted a stalled mitotic state with persistent CCNB:CDK1 activity and cessation of oscillations, mirroring experimental mitotic arrest (**Fig. 11B,C**). We compared doubling times from all analyses, which show increasing doubling times for 2.5 nM with an arrest at 5.0 nM volasertib (**Fig. 11D**).

**Figure 11.**
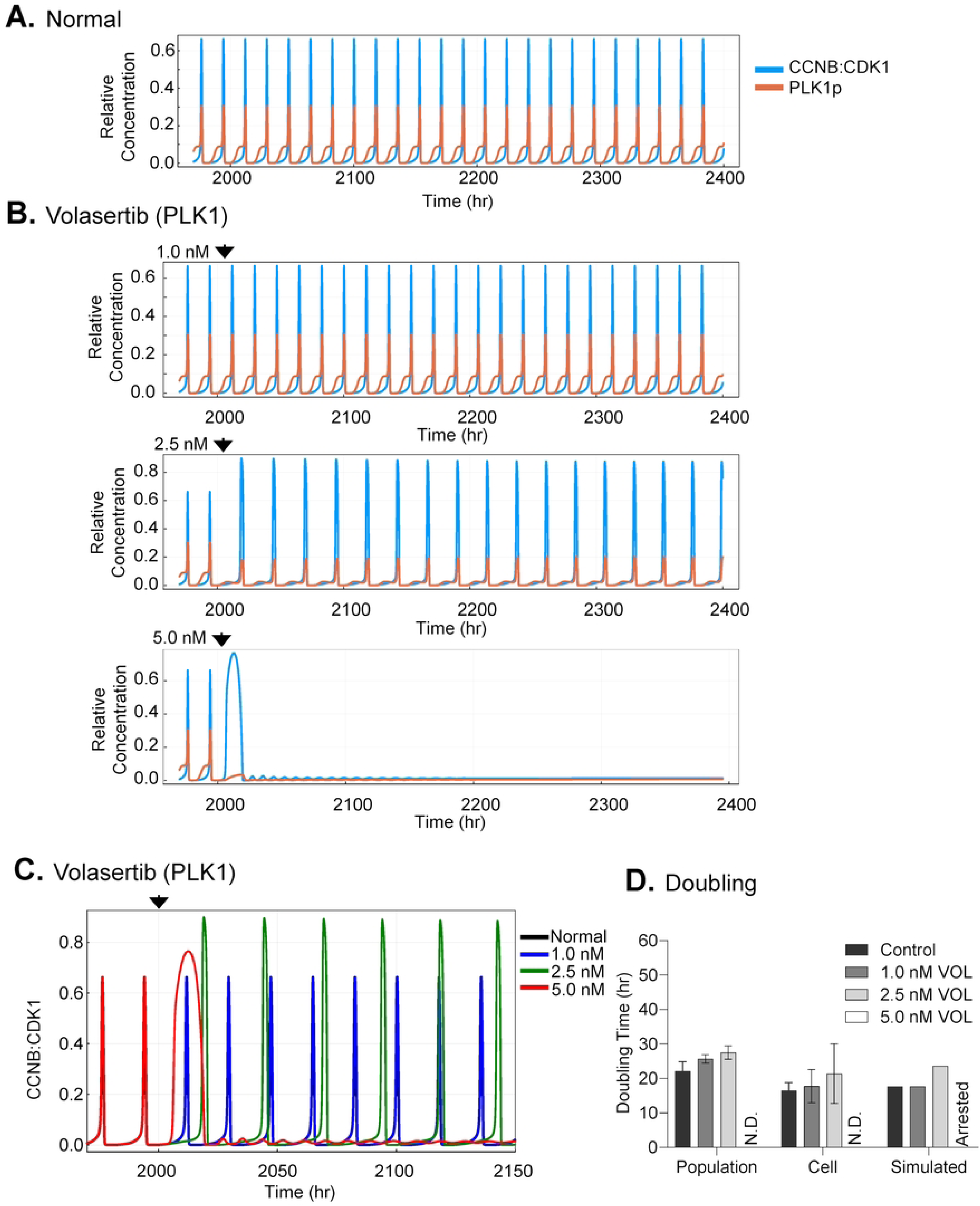
Model responsiveness to volasertib-mediated inhibition of PLK1. **(A)** Baseline simulation of G1 phase regulators, CCNB:CDK1 and PLK1p, under normal conditions, depicting stable, periodic oscillations over time. **(B)** Simulated response to abemaciclib treatment at increasing concentrations (1.0 nM, 2.5 nM, 5.0 nM), demonstrating a dose-dependent slowing of the cell cycle oscillations. As experimentally observed, simulation with 5.0 nM volasertib results in loss of oscillations. **(C)** Overlay of CCNB:CDK1 from (A) and (B). **(D)** Quantitative validation comparing doubling times determined by *in vitro* counting cell numbers of the population, *in vitro* single cell dynamics, and *in silico* simulations under all treatments.

Our overall goal has been to develop a stable, responsive, and oscillatory human cell cycle model grounded in experimental data for *in silico* testing of combinatorial drug effects. To assess combined effects, we analyzed varying combined lower concentrations of abemaciclib (0.001 µM, 0.01 µM, and 0.1 µM) and volasertib (0.4 nM, 1.0 nM, and 2.5 nM) on the proliferation of U251-MG glioma cells (**Fig. 12A**). Cells were treated with DMSO or varying sub-cytotoxic drug combinations, and cell numbers were counted every 24 hrs. Varying abemaciclib with constant 1.0 nM volasertib resulted in an average 30% drop at 0.01 µM abemaciclib and 77% drop at 0.1 µM with little impact of 0.001 µM abemaciclib or volasertib alone after 96 hrs (**Fig. 12A**). Next, we varied volasertib with constant 0.01 µM abemaciclib which resulted in an averaged 47% reduction at 1.0 nM volasertib and 59% at 2.5 nM after 96 hrs (**Fig. 12A**). Addition of 0.01 µM abemaciclib alone resulted in a 22% reduction that was unaffected by addition of 0.4 nM volasertib. To test the predictive power of our model, we simulated these conditions using inhibition parameters calibrated from the single-drug datasets (**Fig. 12B**). Simulated doubling times were converted into population growth curves, enabling direct comparison with experimental measurements. For abemaciclib titrations at 1.0 nM volasertib, we predict cell numbers after 96 hrs using our model to be 1.06 x 10^6^ cells at 0.001 μM abemaciclib (experimental, 1.02 x 10^6^ cells), 1.06 × 10^6^ cells at 0.01 μM (experimental, 0.71 x 10^6^ cells), and 0.51 x 10^6^ cells at 0.1 μM (experimental, 0.23 x 10^6^ cells) (**Fig. 12 A,B**). For volasertib titration at constant 0.01 µM abemaciclib, we simulated 1.36 x 10^6^ cells at 0.4 nM volasertib (experimental, 0.98 x 10^6^ cells), 1.36 x 10^6^ cells at 1.0 nM (experimental, 0.67 x 10^6^ cells), and 0.69 x 10^6^ cells at 2.5 nM (experimentally measured, 0.51 x 10^6^ cells) (**Fig. 12A,B**). Each tested combination differs by at most 2-fold between experimental and simulated data points. Overall, we have developed a holistic, oscillatory mechanistic model of the human cell cycle and have demonstrated that it can simulate the impact of varying concentrations of cell cycle inhibitors comparable to the equivalent experimental *in vitro* data.

**Figure 12.**
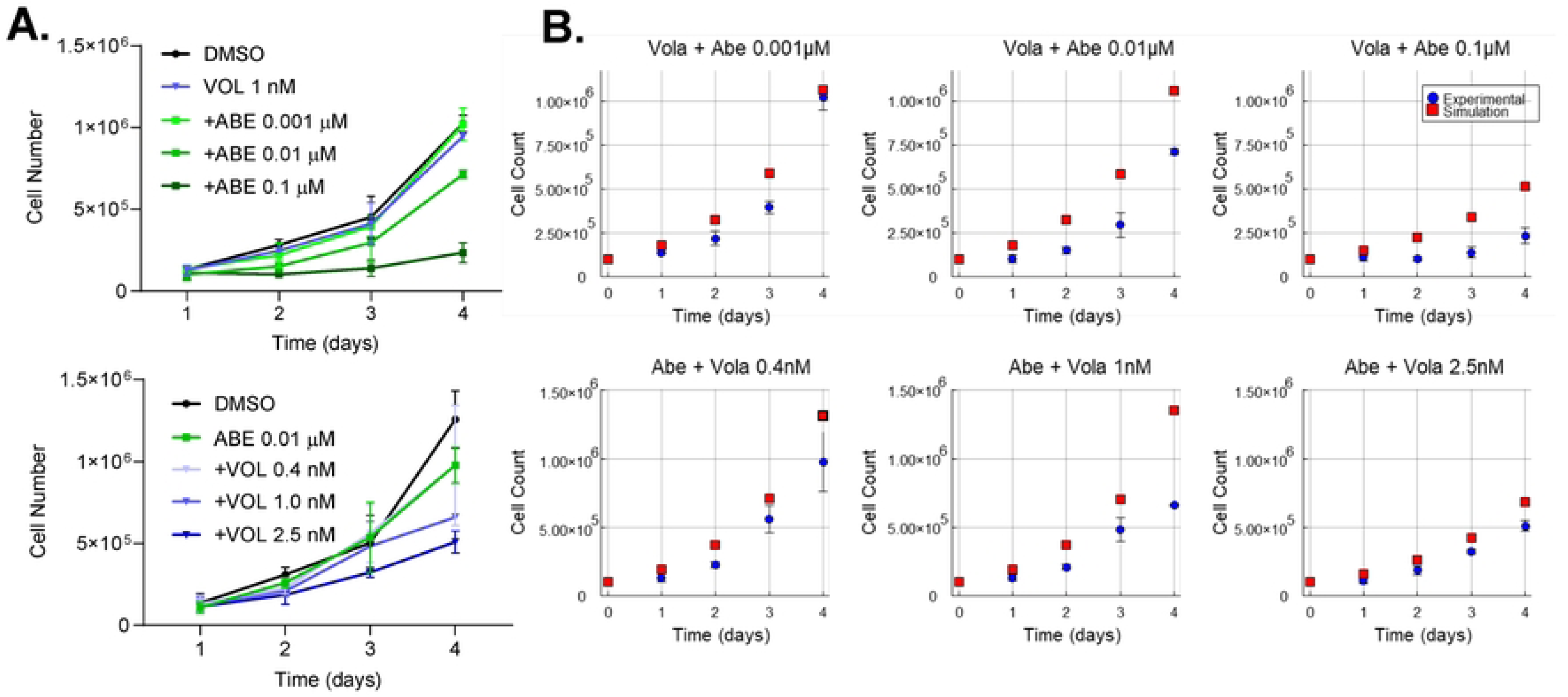
Model validation comparing in silico simulated and in vitro experimental responses to varying concentration to CDK4/6 and PLK1 inhibitors. **(A)** U251-MG FastFUCCI cells were plated subconfluently and treated with the indicated low concentrations of volasertib and abemaciclib or DMSO. Media with inhibitor were replaced every 24 hrs. Cell numbers were determined using a hemocytometer and trypan blue staining out to 96 hrs. Cells were treated with 1 nM volasertib and varying abemaciclib at 0.001 µM, 0.01 µM, 0.1 µM (upper panel), or 0.01 μM abemaciclib and varying volasertib at 0.4 nM, 1.0 nM, 2.5 nM (lower panel). n=3 **(B)** The impact of treatments in (A) on cell cycling were simulated and converted to cell numbers based on the number of oscillations over time for 96 hrs. *In vitro* (blue) and *in silico* (red) results are compared for each condition showing close alignment between simulated and experimental data.

## DISCUSSION

We have developed an experimentally grounded, ordinary differential equation (ODE)-based, comprehensive computational model of the human cell cycle (**Figs. 1 and 2**) to understand network behavior and to test cell cycle inhibitors *in silico*. This new model is predictive and reproduces realistic oscillations and responses to cellular perturbations. Our model was developed by integrating regulatory networks of all four cell cycle phases and calibrating phase timing to the kinetics of the U251 glioma cell line using the FastFUCCI imaging system. Unlike many previous cell cycle models that focus on isolated cellular events or lack quantitative validation, the present model successfully reproduces stable oscillations and accurately predicts the systems-level response to targeted therapeutic perturbations. To build this model, we have expanded upon our previously developed mitosis model [50] to incorporate key G1/S/G2 regulators. The full cell cycle model now includes the activities of non-mitotic cyclin-CDK pairs: CCND:CDK4 (**Fig. 1B**), CCNE:CDK2 (**Fig. 1C**), and CCNA:CDK2 (**Fig. 1D**). It also integrates essential regulatory pathways, including the RB:E2F (**Fig. 2C**), EMI1 **(Fig 2D**), and ATM/Chk2/p21 DNA damage modules (**Fig. 6A**) [51]. Model parameters were defined using a combination of previously published values (directly adopted or fine-tuned) [50, 92, 93] and those estimated from analyses of our cell imaging experiments. It is important to recognize that defining all relationships within a complex network is beyond the ability of any one laboratory. Here, we have used a hybrid framework combining Michaelis-Menten and mass-action kinetics to define the rates of the interacting reactions (see **S2 Appendix** and **S3 Appendix**). The simulations approximate protein-protein interaction behaviors, both direct and indirect, as observed in synchronized human cell cultures and under perturbations that disrupt individual processes. We have constructed this cell cycle model in a modular fashion, enabling future refinements and expansions, as well as the study of disease mechanisms and combinatorial drug therapy.

Our model represents a significant step forward in computational cell cycle modeling by integrating all four phases into a single, unified framework that is both quantitatively calibrated and capable of stable, self-sustaining oscillations. This design addresses major limitations in previous large-scale cell cycle models. For example, the ODE model by Weis et al. [46] required manual resetting of G1 variables to sustain multiple cycles. Similarly, the comprehensive ODE model by Abroudi et al. [93] demonstrated impressive mechanistic scope but produced dampened oscillations and lacked direct quantitative validation. Other large-scale cell cycle models have adopted alternative mathematical formalisms. Other large-scale cell cycle models have adopted alternative mathematical formalisms. The rule-based Lang model [47] achieves remarkable mechanistic depth but relies on a fundamentally different representation of regulatory interactions. Direct comparisons between that framework and our ODE-based kinetic system are therefore challenging. Moreover, as noted by the authors, the Lang model has not yet been experimentally validated for its predicted temporal behaviors. The model also makes several simplifying assumptions, including instantaneous protein synthesis, constant total levels of key regulators, and direct proportionality between Cyclin abundance and Cyclin:CDK complex formation, which may limit its applicability for simulating pharmacological perturbations or cancer-associated dysregulation. Our model overcomes these limitations by maintaining stable, tunable phase timing (**Fig. 7D**) and indefinite oscillations (**Fig. 7E**) while explicitly incorporating the kinetics of protein synthesis, degradation, and regulatory feedback. A key advancement of our work is the alignment of these simulated dynamics with experimental data. We quantitatively parameterized the model using phase durations measured from live-cell FastFUCCI [52] imaging of human U251-MG glioma cells (**Fig. 7A)**Our ODE-based model was specifically designed to simulate pharmacological perturbations by incorporating the kinetics of protein synthesis, degradation, and regulatory feedback, allowing successful validation against inhibitor data. To adapt the model to a new cell type, experimentally measured phase durations and overall doubling time can be used as target constraints for model tuning. Phase-specific timing is adjusted by modifying a small subset of regulatory parameters that govern the G1/S and M/G1 transitions. These adjustments primarily target RB/E2F phosphorylation dynamics and APC/C:CDH1 mediated degradation while preserving the overall network structure. The overall cell cycle length can then be scaled using a global timing parameter (α), allowing the model to match cell-type-specific doubling times without requiring full re-estimation of all parameters. This provides a robust baseline for network perturbation analysis.

### Combinatorial Drug Therapy for Cancer or Viral Pathogenesis

The primary purpose of developing this new integrated computational model is to generate hypotheses and predict new experiments designed to study human disease. Disruptions to the cell cycle network occur in diverse disease states, including cancers [94] and viral infections [95]. In the simplest of terms, cancer is a disease of cell cycle dysfunction defined by loss of checkpoints and development of aneuploidy. The model’s predictive power was demonstrated through simulations of targeted cancer inhibitors. Our simulations of the CDK4/6 inhibitor abemaciclib correctly captured its primary mechanism of action by quantifying how it compresses S/G2/M phases by elongating G1. This culminates in a complete G1 arrest at high concentrations (**Fig. 9B**). While the model accurately predicted the doubling times at medium and high abemaciclib concentrations, it did not fully capture the modest slowing observed at the lowest dose (**Fig. 9D**). In contrast, volasertib simulations were highly consistent with experimental data across all concentrations, faithfully reproducing S/G2/M elongation and mitotic arrest (**Fig. 9D**). Volasertib-induced PLK1 inhibition led to simulated S/G2/M prolongation (+34% at 2.5 nM) and mitotic stalling (**Fig. 9B)**. These simulations matched the experimental phase extension and blebbing, indicating the catastrophe observed in our live-cell imaging experimental studies. We propose that the superior alignment of model predictions with volasertib responses, compared to abemaciclib, reflects the model’s more detailed representation of PLK1 signaling pathways than of CDK4 dynamics. The model’s accuracy in predicting these distinct, phase-specific responses validates the underlying regulatory architecture governing both G1/S progression and mitotic entry.

Analysis of combination treatments showed that dual inhibition resulted in additive suppression of proliferation. Using only the inhibition parameters calibrated from single-drug experiments, the model accurately predicted the combined effects of abemaciclib and volasertib (**Fig 12B)**. Both the experimental data and simulations showed that coupling G1 arrest with mitotic disruption creates a strong block in cycle progression and limits cellular adaptation. The simulated population growth curves under dual inhibition closely matched the experimental data. This agreement shows that the model captures the systems-level interaction between the CDK4/6 and PLK1 pathways. This result is important because it demonstrates that the model can go beyond data fitting and make real predictions about complex drug responses. By using a limited number of experimental data points, we can now predict effective drug concentrations without testing every possible combination *in vitro*. Our findings show how computational models can help prioritize experiments, streamline discovery, and guide the rational design of combination therapies.

### Limitations and Future Directions

While the present model represents a substantial advance in mechanistic simulation of the human cell cycle, several limitations remain that define important directions for future development. The current framework accounts for only a subset of known regulatory interactions, limiting its predictive power for pathways not explicitly represented. For instance, our model does not distinguish between protein family members (e.g., E2F1, E2F2, E2F3, etc.) or make compartmental distinctions between nuclear and cytoplasmic locations. Moreover, because the framework is deterministic, it assumes population uniformity and reflects only average cellular behavior. As such, it does not capture stochastic fluctuations that generate cell-to-cell variability and contribute to heterogeneity in tumor microenvironments. This heterogeneity can be observed in our live cell imaging assays. Additionally, the model has been validated only in U251-MG glioma cells. Extending it to other cell lines or patient-derived models would broaden its applicability. Nonetheless, the strong alignment between simulations and experimental observations supports the validity of the core architecture despite these limitations.

Advancing this model toward broader predictive utility will require progress in three main areas: model refinement, shareability, and applicability. Model refinement will focus on improving regulatory precision and structural realism. While the current framework captures the global logic of cell cycle progression, refinement should incorporate additional molecular detail. Particularly, sub-pathways for drug-specific targets should be represented. For example, many targeted therapies act through downstream cascades such as the PI3K–AKT–mTOR pathway, which influences cyclin stability, checkpoint activation, and overall cell growth [96]. This pathway is currently represented in our model in a simplified form, making it unsuitable for drugs that use this cascade. Similarly, the Aurora kinase B (AURKB) pathway, absent from the present framework, plays a critical role in spindle assembly and cytokinesis [97] and represents an important therapeutic target of drugs such as barasertib [98]. Including such interactions would enable more accurate prediction of therapeutic responses. Similarly, future versions should distinguish between cytoplasmic and nuclear compartments, as compartmentalization governs the localization and timing of key regulators such as CCNB:CDK1, CDC25C, and APC/C. Introducing explicit spatial compartments would improve the mechanistic fidelity of models of processes such as nuclear envelope breakdown and reformation. Model shareability and adaptability are also essential for broader impact. Different cell types exhibit distinct phase durations, checkpoint thresholds, and drug sensitivities. This means that a generalizable modeling framework must allow users to recalibrate it efficiently. Future iterations should include standardized procedures for parameter tuning and guidance for applying the model to other cell lines. In terms of applicability, a key next step toward predictive modeling will be integrating stochastic and population-level dynamics into the current deterministic framework. While the existing ODE system captures average cellular behavior, our FastFUCCI data reveal significant cell-to-cell variability in phase timing, protein levels, and drug response. Reformulating the model using stochastic differential equations (SDEs) or an agent-based approach would enable simulation of single-cell variability and asynchronous population behavior. Coupling this stochastic layer to population-level growth models would bridge molecular regulation and experimental observables such as doubling time, fractional cell death, and drug resistance. Calibration against single-cell and population datasets (such as our FastFUCCI data) will allow prediction of distributions of cellular fates, not just average trends. Such an approach would support *in silico* exploration of treatment responses.

### Summary and Concluding Remarks

Overall, our work establishes a robust, data-driven computational modeling platform for systems-level analysis of the human cell cycle and its disruption by therapeutic perturbations. By integrating experimental data with mechanistic modeling, we demonstrated the power of *in silico* approaches to not only reproduce fundamental cycle dynamics but also forecast responses to targeted inhibitors and their combinations. The model’s modular design, which explicitly incorporates synthesis and degradation of key cell cycle regulators, enables refinement and extension as new mechanisms are needed. This framework is flexible and can be expanded with new pathways, checkpoints, and cell type–specific features. These additions will improve its predictive accuracy. As research on protein-protein interaction networks and regulatory mechanisms advances, the model will remain a useful tool for studying network behavior, investigating disease-related changes, and identifying potential therapeutic targets.

## MATERIALS AND METHODS

### Cell culture and FastFUCCI system

The human glioblastoma multiforme cell line, U251-MG was maintained in DMEM, high glucose (ThermoFisher Scientific) supplemented with 7% fetal bovine serum (FBS) (ThermoFisher Scientific) and 100 U/ml penicillin-streptomycin (ThermoFisher Scientific). Cells were transduced using lentivirus expressing the FastFUCCI system developed by Koh et al [52]. Briefly, the plasmid pBOB-EF1-FastFUCCI-puro (Addgene plasmid #86849) was used to generate a lentiviral vector stock in 293T cells and second-generation packaging plasmids (psPAX2 and pMD2.G). Culture supernatant was collected, filtered, aliquoted and frozen. The U251-MG cells were transduced for 48 hrs then treated with 1 μg/ml puromycin in culture media. The resulting FastFUCCI-labelled U251-MG cells were serial diluted to isolate single colonies enriched for fluorescence.

The FastFUCCI-labelled U251-MG cells were used to measure cell cycle phase timing and impact of cell cycle inhibitors. We used time-lapse imaging with an Incucyte S3 Live-Cell Analysis System (Sartorius) set at 20x magnification, well area 876.2 mm², green acquisition time 200 ms, and red acquisition time 400 ms to capture higher-resolution proliferation dynamics. Cells were plated in six-well plates at 1 x 10^4^ cells per well, then serum-starved using DMEM supplemented 0.1% FBS with penicillin-streptomycin for 48 hr. Subsequently, the media was replaced with DMEM containing 7% FBS and penicillin-streptomycin, then imaged every 2 hrs to 96 hrs, collecting 16 fields of view per well. Images were exported as both still images and movies, and we manually tracked cells using still images (e.g., cell #1 to #1.1 and #1.2, then cell #1.1 to #1.1.1 and #1.1.2, etc.). Mitotic events were annotated by visualizing one rounded, geminin-green expressing cell becoming two daughter cells lacking the signal. We defined cell doubling time as the interval between successive mitoses in the same tracked cell (from the first to the second division). Cell cycle phase durations were recorded using the FastFUCCI reporter system [52], in which nuclei fluoresce red during G1 phase and green during S/G2/M phases, enabling extraction of phase-specific timings from single-cell traces. Doubling times and phase durations were averaged across tracked cells, with standard deviations reported.

Cells were treated with inhibitors, abemaciclib, selective against CDK4 (Selleck Chemicals #55716), volasertib, selective against PLK1 (Selleck Chemicals #52235) or DMSO (Sigma-Aldrich #D8418). Compounds were resuspended in DMSO as directed by the manufacturer. Cells were plated as above and treated with abemaciclib at 0.01 μM, 0.1 μM, or 1.0 μM, or volasertib at 1 nM, 2.5 nM, or 5 nM. We quantified changes in cell proliferation using both IncuCyte imaging and hemocytometer counting. Cultures used for counting had compounds changed every 24 hrs, while IncuCyte imaging went undisrupted for the duration of the time. Cell counts from replicate wells were averaged to generate growth curves for each treatment group. Dual-drug combination studies were done by holding one drug at a constant concentration while applying various concentrations of the other. Cells were initially plated at 1 x 10^5^ cells per well. Media was changed every day with appropriate drug concentrations and cells numbers determined using a hemocytometer. Assuming exponential population growth, the cell number as a function of time was modeled as

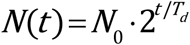

where N(t) is the number of cells at time t, N_0_ is the initial number of cells, and T_d_ is the doubling time. Linearization was performed by applying the base-2 logarithm:

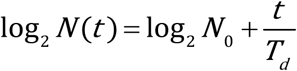

A linear model of the form y=mx+b was then fit to the transformed data, with y=log_2_N and x = t. The slope of the fitted line, m, corresponds to 1/T_d_. Thus, the doubling time was calculated as:

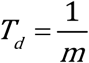

Doubling times were reported in days and converted to hours by multiplying by 24. Fitting was restricted to time points prior to confluence or nutrient depletion to ensure exponential growth dynamics. Doubling times were averaged across replicates, with standard deviations calculated to assess variability.

### Model Development

Based on prior biological knowledge of cell cycle regulation and our established molecular network of the mitotic cell cycle, we constructed a mechanistic biochemical network comprising 63 state variables, representing unphosphorylated proteins, phosphorylated proteins, and protein complexes (**Fig. 1 and 2**). Protein nomenclature follows UniProt standards, with a suffix “P” denoting phosphorylated forms. The network includes 41 major biochemical reactions, encompassing synthesis, degradation, phosphorylation, dephosphorylation, association, and dissociation processes among core cell cycle regulators (detailed in **S2 Appendix**). We note that the cell cycle biopathway does not distinguish between reactions occurring within the cytosolic and nucleus compartments. The model was initially constructed using BioModME [99] to ensure the accuracy of the equations and to provide programming language-level code for further analysis.

The biochemical network was formalized as a dynamic computational model consisting of 63 ordinary differential equations (ODEs) describing the temporal dynamics of protein concentrations (normalized to total CDK1 concentration, denoted [X] for protein X; **S3 Appendix**). Reaction rates were modeled using a hybrid formulation of mass action and Michaelis–Menten kinetics. The model comprises 218 kinetic, synthesis, and degradation parameters (**S4 Appendix**). Protein synthesis is described by rate constants k_s_ and degradation rate of a protein through k_d_. Forward rate constants (k_f_) represent cycle-promoting events such as phosphorylation or complex formation, and reverse rate constants (k_r_) represent antagonistic reactions such as dephosphorylation or dissociation. Degradation processes occur via multiple routes, including self-degradation and targeted degradation mediated by APC/CP:CDC20, APC/C:CDH1, and SCF, as established experimentally.

Drug mechanisms of action were incorporated through Hill-type inhibition factors that multiplicatively modulated phosphorylation and activation terms in the relevant reactions. For a given drug, the inhibition factor is defined as

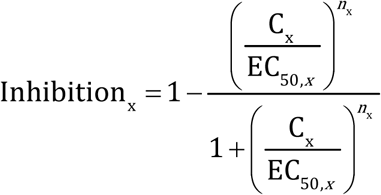

where C_x_ is the drug concentration, EC_50,x_ is the half-maximal effective concentration, and n_x_ is the Hill coefficient. For abemaciclib, this inhibition factor was applied to CDK4 activity; for volasertib, it was applied to PLK1 activity.

### Model Simulations

We solved the 63 ordinary differential equations (ODEs) listed in **S3 Appendix** with parameter values from **S4 Appendix** using Julia’s AutoTsit5–Rosenbrock23 adaptive solver, which switches between 5th-order Runge-Kutta and 3rd-order Rosenbrock methods based on system stiffness. Simulations were performed in Julia (version 1.11.6) with relative and absolute tolerances set to (10^-6^). Results are presented in **Figs. 3–12**. The Julia code is available at https://github.com/MCWComputationalBiologyLab/Womack_CellCycle_2025 with equivalent MATLAB (version R2024b) code provided for reproducibility.

Callback functions in Julia were used to implement dynamic events by applying parameter changes at specific time points. For mitogen withdrawal and re-addition (**Fig. 5**), the synthesis rate of cyclin D (ks_CCND_) was toggled between baseline values and starved conditions (ks_CCND_=0). For DNA damage (**Fig. 6**), a pulse was modeled by setting the ATM phosphorylation rate (kf_ATMp_) to 50 at the onset of damage and to 0 after sustained damage, with no-damage baselines used otherwise. For drug treatments (**Fig. 9 and 11**), concentrations of abemaciclib (0.01, 0.1, 1.0 μM) and volasertib (1.0, 2.5, 5.0 nM) were applied at specific time points (e.g., volasertib at 1.0 nM at t = 2000 hours) as model parameters. The timescale and cell cycle period were controlled by a scaling factor (α), tuned to 2.3 based on experimental data to yield a cell cycle period of 17.73 hours.

Cell cycle progression was monitored by defining phase transitions based on molecular markers and regulatory events. The G1/S transition was marked by the maximal concentration of cyclin E [76], S/G2 by half-maximal crossings of phosphorylated PLK1 [67, 77], G2/M by local maxima of phosphorylated CDC25C [78], and M/G1 by the sharp post-mitotic decline in phosphorylated lamin A/C (LMNAp). These markers enabled annotation of individual cell cycle phases, calculation of relative phase durations, and validation against experimental doubling times from cell counting experiments.

### Model Parameterization

The mitotic cell cycle model was initialized with a previously established set of 105 parameter values [50]. These were adjusted to ensure consistent coupling between mitotic events and other cell cycle phases. Additional parameters were iteratively incorporated, starting with a simplified cycle model and progressively adding regulatory details for feedbacks and checkpoints.

To ensure biologically realistic dynamics, candidate parameter sets were screened using a constraint-based framework. Simulations were evaluated against four criteria: (1) sustained oscillations with consistent period and amplitude over 10,000 hours, (2) correct ordering of cell cycle events, with PLK1 activation preceding mitotic entry (defined by the rise of CCNB:CDK1, also termed mitosis-promoting factor) and sequential activation of cyclin–CDK complexes (CCND:CDK4, CCNE:CDK2, CCNA:CDK2, CCNB:CDK1), (3) temporal allocation of phases matching experimental data for U251-MG cells, with G1 comprising approximately 60% of the cell cycle and mitosis (CCNB:CDK1 activity) less than 10% of the doubling time, and (4) maximal activities of cyclin–CDK complexes constrained within the same order of magnitude to prevent unrealistic disparities. With these criteria established, parameter exploration was performed using a user-defined set anchored to baseline values. For each parameter, ranges were specified through fixed percentage variations or explicit bounds, and the Cartesian product of these ranges was employed to construct the complete parameter grid. Simulations were performed for each parameter set, and results that satisfied all biological checks were recorded and curated for downstream analysis.

Drug-specific parameters for abemaciclib and volasertib were estimated by fitting the Hill-type inhibition functions to experimental dose–response data. Half-maximal effective concentrations (EC50) and Hill coefficients were optimized to minimize the least-squares error between simulated outputs (doubling times and phase durations) and experimental datasets. Optimization was performed using the global nonlinear optimization package Optim.jl in Julia. The final parameter values reported in **S4 Appendix**, correspond to the best-fit solutions that reproduced experimental readouts.

## ACKNOWLEDGEMENTS

We thank members of the Terhune and Dash laboratories, along with staff in the Department of Biomedical Engineering, for their assistance. We thank the MCW Center for Immunology for the use of the Incucyte instrument. Figure 1A was created in BioRender. This work was supported by a grant from Advancing a Healthier Wisconsin Endowment to S.S.T. and R.K.D. and the Jeschke Scholarship Fund from the Medical College of Wisconsin to A.J.F. Additional support was provided by a generous philanthropic gift from The Stead Family Foundation. The funders had no role in study design, data collection and analysis, decision to publish, or preparation of the manuscript.

## SUPPORTING INFORMATION

**S1 Appendix.** The gene and protein names involved in the human cell cycle system.

**S2 Appendix.** The major biochemical reactions and other related reactions among the mitotic proteins and the associated protein complexes, and their descriptions based on the mitotic biopathway of **Figs. 1 and 2**.

**S3 Appendix.** The ODEs (ordinary differential equations) that describe the dynamics of the cell cycle proteins and the associated protein complexes.

**S4 Appendix.** Table of model parameters.

## AUTHOR CONTRIBUTIONS

**Conceptualization**: J.A.W., S.S.T., and R.K.D. **Data curation:** A.T.S., A.J.F., and J.A.W. **Formal analysis:** J.A.W., S.S.T., and R.K.D. **Funding acquisition:** .S.S.T. and R.K.D. **Investigation:** A.T.S., A.J.F., and J.A.W. **Methodology:** J.A.W., S.S.T., and R.K.D. **Supervision:** S.S.T., R.K.D. **Validation:** J.A.W., S.S.T., and R.K.D. **Visualization**: J.A.W. and S.S.T., **Writing – original draft:** A.T.S., A.J.F., J.A.W., S.S.T. and R.K.D. **Writing – review & editing**: J.A.W., S.S.T., and R.K.D.

## Notes

### Competing Interest Statement

The authors have declared no competing interest.

## References

1. Park MS, Koff A. Overview of the cell cycle. Current protocols in cell biology. 1998;(1):8.1.-8.1. 9.

2. Pennycook BR, Barr AR. Restriction point regulation at the crossroads between quiescence and cell proliferation. FEBS letters. 2020;594(13):2046–60.

3. Hanahan D. Hallmarks of cancer: new dimensions. Cancer discovery. 2022;12(1):31–46.

4. Molinari M. Cell cycle checkpoints and their inactivation in human cancer. Cell proliferation. 2000;33(5):261–74.

5. Vermeulen K, Van Bockstaele DR, Berneman ZN. The cell cycle: a review of regulation, deregulation and therapeutic targets in cancer. Cell Proliferation. 2003;36(3):131–49. doi: 10.1046/j.1365-2184.2003.00266.x.

6. Lapenna S, Giordano A. Cell cycle kinases as therapeutic targets for cancer. Nature reviews Drug discovery. 2009;8(7):547–66.

7. Malumbres M, Barbacid M. Cell cycle, CDKs and cancer: a changing paradigm. Nature Reviews Cancer. 2009;9:153. doi: 10.1038/nrc2602.

8. Wang Z. Regulation of cell cycle progression by growth factor-induced cell signaling. Cells. 2021;10(12):3327.

9. Cheung TH, Rando TA. Molecular regulation of stem cell quiescence. Nature reviews Molecular cell biology. 2013;14(6):329–40.

10. Foster DA, Yellen P, Xu L, Saqcena M. Regulation of G1 cell cycle progression: distinguishing the restriction point from a nutrient-sensing cell growth checkpoint (s). Genes & cancer. 2010;1(11):1124–31.

11. Davey NE, Morgan DO. Building a regulatory network with short linear sequence motifs: lessons from the degrons of the anaphase-promoting complex. Molecular cell. 2016;64(1):12–23.

12. Pines J. Cubism and the cell cycle: the many faces of the APC/C. Nature reviews Molecular cell biology. 2011;12(7):427–38.

13. Lindqvist A, Rodríguez-Bravo V, Medema RH. The decision to enter mitosis: feedback and redundancy in the mitotic entry network. Journal of Cell Biology. 2009;185(2):193–202.

14. Malumbres M. Cyclin-dependent kinases. Genome biology. 2014;15(6):122.

15. Verdugo A, Vinod PK, Tyson JJ, Novak B. Molecular mechanisms creating bistable switches at cell cycle transitions. Open Biol. 2013;3(3):120179. doi: 10.1098/rsob.120179. PubMed PMID: 23486222; PubMed Central PMCID: PMCPMC3718337.

16. Stallaert W, Kedziora KM, Chao HX, Purvis JE. Bistable switches as integrators and actuators during cell cycle progression. FEBS letters. 2019;593(20):2805–16.

17. Choudhury R, Bonacci T, Arceci A, Lahiri D, Mills CA, Kernan JL, et al. APC/C and SCFcyclin F constitute a reciprocal feedback circuit controlling S-phase entry. Cell reports. 2016;16(12):3359–72.

18. Satyanarayana A, Kaldis P. Mammalian cell-cycle regulation: several Cdks, numerous cyclins and diverse compensatory mechanisms. Oncogene. 2009;28(33):2925–39. doi: 10.1038/onc.2009.170.

19. Besson A, Dowdy SF, Roberts JM. CDK inhibitors: cell cycle regulators and beyond. Developmental cell. 2008;14(2):159–69.

20. Lim S, Kaldis P. Cdks, cyclins and CKIs: roles beyond cell cycle regulation. Development. 2013;140(15):3079–93.

21. Mudryj M, Devoto SH, Hiebert SW, Hunter T, Pines J, Nevins JR. Cell cycle regulation of the E2F transcription factor involves an interaction with cyclin A. Cell. 1991;65(7):1243–53.

22. Geng Y, Eaton EN, Picon M, Roberts JM, Lundberg AS, Gifford A, et al. Regulation of cyclin E transcription by E2Fs and retinoblastoma protein. Oncogene. 1996;12(6):1173–80.

23. Lukas C, Sorensen CS, Kramer E, Santoni-Rugiu E, Lindeneg C, Peters JM, et al. Accumulation of cyclin B1 requires E2F and cyclin-A-dependent rearrangement of the anaphase-promoting complex. Nature. 1999;401(6755):815–8. Epub 1999/11/05. doi: 10.1038/44611. PubMed PMID: 10548110.

24. Stevens C, La Thangue NB. E2F and cell cycle control: a double-edged sword. Archives of biochemistry and biophysics. 2003;412(2):157–69.

25. Zachariae W, Schwab M, Nasmyth K, Seufert W. Control of Cyclin Ubiquitination by CDK-Regulated Binding of Hct1 to the Anaphase Promoting Complex. Science. 1998;282(5394):1721.

26. Kramer ER, Scheuringer N, Podtelejnikov AV, Mann M, Peters J-M. Mitotic regulation of the APC activator proteins CDC20 and CDH1. Molecular biology of the cell. 2000;11(5):1555–69.

27. Ciliberto A, Lukács A, Tóth A, TysonÍ JJ, Novák B. Rewiring the exit from mitosis. Cell Cycle. 2005;4(8):4107–12.

28. Castro A, Bernis C, Vigneron S, Labbe J-C, Lorca T. The anaphase-promoting complex: a key factor in the regulation of cell cycle. Oncogene. 2005;24(3):314–25.

29. Amador V, Ge S, Santamaría PG, Guardavaccaro D, Pagano M. APC/CCdc20 controls the ubiquitin-mediated degradation of p21 in prometaphase. Molecular cell. 2007;27(3):462–73.

30. Pesin JA, Orr-Weaver TL. Regulation of APC/C activators in mitosis and meiosis. Annual review of cell and developmental biology. 2008;24(1):475–99.

31. Li M, Zhang P. The function of APC/C Cdh1 in cell cycle and beyond. Cell division. 2009;4:1–7.

32. Koepp DM, Schaefer LK, Ye X, Keyomarsi K, Chu C, Harper JW, et al. Phosphorylation-dependent ubiquitination of cyclin E by the SCFFbw7 ubiquitin ligase. Science. 2001;294(5540):173–7.

33. Ang XL, Wade Harper J. SCF-mediated protein degradation and cell cycle control. Oncogene. 2005;24(17):2860–70. Epub 2005/04/20. doi: 10.1038/sj.onc.1208614. PubMed PMID: 15838520.

34. Thompson LL, Rutherford KA, Lepage CC, McManus KJ. The SCF complex is essential to maintain genome and chromosome stability. International Journal of Molecular Sciences. 2021;22(16):8544.

35. Health NIo. NIH to prioritize human-based research technologies Bethesda, MD: National Institutes of Health; 2025 [updated April 29 2025; cited 2025 November 1 2025]. Available from: https://www.nih.gov/news-events/news-releases/nih-prioritize-human-based-research-technologies.

36. Tyson JJ, Novák B. Time-keeping and decision-making in the cell cycle. Interface Focus. 2022;12(4):20210075.

37. Novak B, Tyson JJ. A model for restriction point control of the mammalian cell cycle. J Theor Biol. 2004;230(4):563–79. doi: 10.1016/j.jtbi.2004.04.039. PubMed PMID: 15363676.

38. Aguda B, Tang Y. The kinetic origins of the restriction point in the mammalian cell cycle. Cell proliferation. 1999;32(5):321–35.

39. Aguda BD. A quantitative analysis of the kinetics of the G2 DNA damage checkpoint system. Proceedings of the National Academy of Sciences. 1999;96(20):11352–7.

40. Tyson JJ. Modeling the cell division cycle: cdc2 and cyclin interactions. Proceedings of the National Academy of Sciences. 1991;88(16):7328–32. doi: 10.1073/pnas.88.16.7328.

41. Obeyesekere M, Knudsen E, Wang J, Zimmerman S. A mathematical model of the regulation of the G1 phase of Rb+/+ and Rb—/—mouse embryonic fibroblasts and an osteosarcoma cell line. Cell proliferation. 1997;30(3-4):171–94.

42. Kohn KW. Functional capabilities of molecular network components controlling the mammalian G1/S cell cycle phase transition. Oncogene. 1998;16(8):1065–75.

43. Hatzimanikatis V, Lee K, Bailey J. A mathematical description of regulation of the G1-S transition of the mammalian cell cycle. Biotechnology and bioengineering. 1999;65(6):631–7.

44. Obeyesekere MN, Zimmerman SO, Tecarro ES, Auchmuty G. A model of cell cycle behavior dominated by kinetics of a pathway stimulated by growth factors. Bulletin of mathematical biology. 1999;61(5):917–34.

45. Ling H, Kulasiri D, Samarasinghe S. Robustness of G1/S checkpoint pathways in cell cycle regulation based on probability of DNA-damaged cells passing as healthy cells. Biosystems. 2010;101(3):213–21.

46. Weis MC, Avva J, Jacobberger JW, Sreenath SN. A data-driven, mathematical model of mammalian cell cycle regulation. PLoS One. 2014;9(5):e97130.

47. Lang PF, Penas DR, Banga JR, Weindl D, Novak B. Reusable rule-based cell cycle model explains compartment-resolved dynamics of 16 observables in RPE-1 cells. PLoS computational biology. 2024;20(1):e1011151.

48. Gérard C, Goldbeter A. A skeleton model for the network of cyclin-dependent kinases driving the mammalian cell cycle. Interface Focus. 2011;1(1):24–35.

49. Gerard C, Goldbeter A. Temporal self-organization of the cyclin/Cdk network driving the mammalian cell cycle. Proceedings of the National Academy of Sciences. 2009;106(51):21643–8. doi: 10.1073/pnas.0903827106.

50. Terhune SS, Jung Y, Cataldo KM, Dash RK. Network mechanisms and dysfunction within an integrated computational model of progression through mitosis in the human cell cycle. PLoS computational biology. 2020;16(4):e1007733.

51. Jung Y, Kraikivski P, Shafiekhani S, Terhune SS, Dash RK. Crosstalk between Plk1, p53, cell cycle, and G2/M DNA damage checkpoint regulation in cancer: computational modeling and analysis. npj Systems Biology and Applications. 2021;7(1):46.

52. Koh S-B, Mascalchi P, Rodriguez E, Lin Y, Jodrell DI, Richards FM, et al. A quantitative FastFUCCI assay defines cell cycle dynamics at a single-cell level. Journal of cell science. 2017;130(2):512–20.

53. Cataldo KM, Roche KL, Monti CE, Dash RK, Murphy EA, Terhune SS. The effective multiplicity of infection for HCMV depends on the activity of the cellular 20S proteasome. Journal of Virology. 2025;99(1):e01751–24.

54. Winston JT, Pledger W. Growth factor regulation of cyclin D1 mRNA expression through protein synthesis-dependent and-independent mechanisms. Molecular biology of the cell. 1993;4(11):1133–44.

55. Lavoie JN, L’Allemain G, Brunet A, Müller R, Pouysségur J. Cyclin D1 expression is regulated positively by the p42/p44MAPK and negatively by the p38/HOGMAPK pathway. Journal of Biological Chemistry. 1996;271(34):20608–16.

56. Muise-Helmericks RC, Grimes HL, Bellacosa A, Malstrom SE, Tsichlis PN, Rosen N. Cyclin D expression is controlled post-transcriptionally via a phosphatidylinositol 3-kinase/Akt-dependent pathway. Journal of Biological Chemistry. 1998;273(45):29864–72.

57. Kato J-Y, Matsuoka M, Strom DK, Sherr CJ. Regulation of cyclin D-dependent kinase 4 (cdk4) by cdk4-activating kinase. Molecular and cellular biology. 1994;14(4):2713–21.

58. Narasimha AM, Kaulich M, Shapiro GS, Choi YJ, Sicinski P, Dowdy SF. Cyclin D activates the Rb tumor suppressor by mono-phosphorylation. Elife. 2014;3:e02872.

59. Tuck C, Zhang T, Potapova T, Malumbres M, Novák B. Robust mitotic entry is ensured by a latching switch. Biology open. 2013;2(9):924–31.

60. Carrano AC, Eytan E, Hershko A, Pagano M. SKP2 is required for ubiquitin-mediated degradation of the CDK inhibitor p27. Nature cell biology. 1999;1(4):193–9.

61. Nakayama K, Nagahama H, Minamishima YA, Matsumoto M, Nakamichi I, Kitagawa K, et al. Targeted disruption of Skp2 results in accumulation of cyclin E and p27Kip1, polyploidy and centrosome overduplication. The EMBO journal. 2000.

62. Pagano M, Pepperkok R, Verde F, Ansorge W, Draetta G. Cyclin A is required at two points in the human cell cycle. The EMBO journal. 1992;11(3):961–71.

63. Gu Y, Rosenblatt J, Morgan D. Cell cycle regulation of CDK2 activity by phosphorylation of Thr160 and Tyr15. The EMBO journal. 1992;11(11):3995–4005.

64. Gabrielli BG, Lee MS, Walker DH, Piwnica-Worms H, Maller JL. Cdc25 regulates the phosphorylation and activity of the Xenopus cdk2 protein kinase complex. Journal of Biological Chemistry. 1992;267(25):18040–6.

65. Sur S, Agrawal DK. Phosphatases and kinases regulating CDC25 activity in the cell cycle: clinical implications of CDC25 overexpression and potential treatment strategies. Molecular and cellular biochemistry. 2016;416:33–46.

66. Watanabe N, Arai H, Nishihara Y, Taniguchi M, Watanabe N, Hunter T, et al. M-phase kinases induce phospho-dependent ubiquitination of somatic Wee1 by SCFβ-TrCP. Proceedings of the National Academy of Sciences. 2004;101(13):4419–24.

67. Cascales HS, Burdova K, Middleton A, Kuzin V, Müllers E, Stoy H, et al. Cyclin A2 localises in the cytoplasm at the S/G2 transition to activate PLK1. Life Science Alliance. 2021;4(3).

68. Kalous J, Aleshkina D. Multiple roles of PLK1 in mitosis and meiosis. Cells. 2023;12(1):187.

69. Lindon C, Pines J. Ordered proteolysis in anaphase inactivates Plk1 to contribute to proper mitotic exit in human cells. Journal of Cell Biology. 2004;164(2):233–41.

70. Mittnacht S. Control of pRB phosphorylation. Current opinion in genetics & development. 1998;8(1):21–7.

71. Hsu JY, Reimann JD, Sørensen CS, Lukas J, Jackson PK. E2F-dependent accumulation of hEmi1 regulates S phase entry by inhibiting APCCdh1. Nature cell biology. 2002;4(5):358–66.

72. Qiao R, Weissmann F, Yamaguchi M, Brown NG, VanderLinden R, Imre R, et al. Mechanism of APC/CCDC20 activation by mitotic phosphorylation. Proceedings of the National Academy of Sciences. 2016;113(19):E2570–E8.

73. Rossio V, Michimoto T, Sasaki T, Ohbayashi I, Kikuchi Y, Yoshida S. Nuclear PP2A-Cdc55 prevents APC-Cdc20 activation during the spindle assembly checkpoint. Journal of cell science. 2013;126(19):4396–405.

74. Hein JB, Hertz EP, Garvanska DH, Kruse T, Nilsson J. Distinct kinetics of serine and threonine dephosphorylation are essential for mitosis. Nature cell biology. 2017;19(12):1433–40.

75. Robbins JA, Cross FR. Regulated degradation of the APC coactivator Cdc20. Cell division. 2010;5:1–9.

76. Clurman BE, Sheaff RJ, Thress K, Groudine M, Roberts JM. Turnover of cyclin E by the ubiquitin-proteasome pathway is regulated by cdk2 binding and cyclin phosphorylation. Genes & development. 1996;10(16):1979–90.

77. Barr AR, Heldt FS, Zhang T, Bakal C, Novak B. A dynamical framework for the all-or-none G1/S transition. Cell systems. 2016;2(1):27–37.

78. Lemonnier T, Dupré A, Jessus C. The G2-to-M transition from a phosphatase perspective: a new vision of the meiotic division. Cell Division. 2020;15(1):9.

79. Chehab NH, Malikzay A, Appel M, Halazonetis TD. Chk2/hCds1 functions as a DNA damage checkpoint in G1 by stabilizing p53. Genes & development. 2000;14(3):278–88.

80. Elledge SJ. Cell cycle checkpoints: preventing an identity crisis. Science. 1996;274(5293):1664–72.

81. Chen L, Gilkes DM, Pan Y, Lane WS, Chen J. ATM and Chk2-dependent phosphorylation of MDMX contribute to p53 activation after DNA damage. The EMBO journal. 2005;24(19):3411–22.

82. Barak Y, Juven T, Haffner R, Oren M. mdm2 expression is induced by wild type p53 activity. The EMBO journal. 1993;12(2):461–8.

83. Harper JW, Adami GR, Wei N, Keyomarsi K, Elledge SJ. The p21 Cdk-interacting protein Cip1 is a potent inhibitor of G1 cyclin-dependent kinases. Cell. 1993;75(4):805–16.

84. Adkins JN, Lumb KJ. Stoichiometry of Cyclin A− Cyclin-Dependent Kinase 2 Inhibition by p21Cip1/Waf1. Biochemistry. 2000;39(45):13925–30.

85. Hengst L, Göpfert U, Lashuel HA, Reed SI. Complete inhibition of Cdk/cyclin by one molecule of p21Cip1. Genes & development. 1998;12(24):3882–8.

86. Harper JW, Elledge SJ, Keyomarsi K, Dynlacht B, Tsai L-H, Zhang P, et al. Inhibition of cyclin-dependent kinases by p21. Molecular biology of the cell. 1995;6(4):387–400.

87. Ballabeni A, Zamponi R, Moore JK, Helin K, Kirschner MW. Geminin deploys multiple mechanisms to regulate Cdt1 before cell division thus ensuring the proper execution of DNA replication. Proceedings of the National Academy of Sciences. 2013;110(30):E2848–E53.

88. Kim ES. Abemaciclib: first global approval. Drugs. 2017;77:2063–70.

89. Poratti M, Marzaro G. Third-generation CDK inhibitors: A review on the synthesis and binding modes of Palbociclib, Ribociclib and Abemaciclib. European Journal of Medicinal Chemistry. 2019;172:143–53.

90. Martin JM, Goldstein LJ. Profile of abemaciclib and its potential in the treatment of breast cancer. OncoTargets and therapy. 2018:5253–9.

91. Gjertsen BT, Schöffski P. Discovery and development of the Polo-like kinase inhibitor volasertib in cancer therapy. Leukemia. 2015;29(1):11–9.

92. Tyson JJ, Novak B. Temporal organization of the cell cycle. Current biology : CB. 2008;18(17):R759-r68. Epub 2008/09/13. doi: 10.1016/j.cub.2008.07.001. PubMed PMID: 18786381; PubMed Central PMCID: PMCPMC2856080.

93. Abroudi A, Samarasinghe S, Kulasiri D. A comprehensive complex systems approach to the study and analysis of mammalian cell cycle control system in the presence of DNA damage stress. Journal of theoretical biology. 2017;429:204–28.

94. Matthews HK, Bertoli C, de Bruin RA. Cell cycle control in cancer. Nature Reviews Molecular Cell Biology. 2022;23(1):74–88.

95. Fan Y, Sanyal S, Bruzzone R. Breaking bad: how viruses subvert the cell cycle. Frontiers in cellular and infection microbiology. 2018;8:396.

96. Gao N, Zhang Z, Jiang B-H, Shi X. Role of PI3K/AKT/mTOR signaling in the cell cycle progression of human prostate cancer. Biochemical and biophysical research communications. 2003;310(4):1124–32.

97. Ducat D, Zheng Y. Aurora kinases in spindle assembly and chromosome segregation. Experimental cell research. 2004;301(1):60–7.

98. Helfrich BA, Kim J, Gao D, Chan DC, Zhang Z, Tan A-C, et al. Barasertib (AZD1152), a small molecule Aurora B inhibitor, inhibits the growth of SCLC cell lines in vitro and in vivo. Molecular cancer therapeutics. 2016;15(10):2314-22.

99. Womack JA, Shah V, Audi SH, Terhune SS, Dash RK. BioModME for building and simulating dynamic computational models of complex biological systems. Bioinformatics Advances. 2024;4(1):vbae023.

